# Architecture of the MKK6-p38α complex defines the basis of MAPK specificity and activation

**DOI:** 10.1101/2022.07.04.498667

**Authors:** Pauline Juyoux, Ioannis Galdadas, Dorothea Gobbo, Jill von Velsen, Martin Pelosse, Mark Tully, Oscar Vadas, Francesco Luigi Gervasio, Erika Pellegrini, Matthew W. Bowler

## Abstract

The MAP kinase p38α is a central component of signalling in inflammation and the immune response and is, therefore, an important drug target. Little is known about the molecular mechanism of its activation by double-phosphorylation from MAP2Ks, due to the challenge of trapping a transient and dynamic hetero-kinase complex. Here, we applied a multidisciplinary approach to generate the first structure of p38α in complex with its MAP2K MKK6 and understand the activation mechanism. Integrating cryo-EM with MD simulations, HDX-MS and *in cellulo* experiments, we demonstrate a dynamic, multi-step, phosphorylation mechanism, reveal new catalytically relevant interactions, and show that MAP2K disordered N-termini determine pathway specificity. Our work captures, for the first time, a fundamental step of cell signalling: a kinase phosphorylating its downstream target kinase.

**One-Sentence Summary:** Integrative Cryo-EM and MD analysis of an active hetero-kinase complex reveals details of cellular signal transmission

## Main Text

The mitogen activated protein kinases (MAPKs) are conserved in all eukaryotes where they form signalling cascades responding to extra-cellular stimuli leading to diverse responses from differentiation to apoptosis. In higher organisms, the MAPK p38α acts in response to stress such as irradiation, hypoxia and osmotic shock, and also to signalling from inflammatory cytokines (*1*). Pathogens, including SARS-CoV-2, often elicit upregulation of p38α, which can lead to the cytokine storm associated with severe COVID-19 (*2, 3*). Signals propagate through phosphorylation of successive protein kinases (MAP4K, MAP3K to MAP2K), which eventually phosphorylate and activate a MAPK via double phosphorylation at the TxY motif in the activation loop (A-loop), leading to a conformational change (*4, 5*). When the A-loop is phosphorylated, key residues surrounding the ATP and Mg^2+^ ions reorient, stabilising the activated MAPK. The activated MAPK is then transported to the nucleus where it modulates gene expression (*6–8*). The key role of p38α in inflammation, and the fact that aberrant p38α signalling is implicated in numerous diseases, such as arthritis and cancer, but also in the response to infection, make it a highly studied drug target (*1*). Despite initial successes in developing potent compounds targeting the kinase nucleotide binding pocket, many potential therapies failed in clinical trials due to off-target effects (*9*). Therefore, a molecular understanding of its interaction with upstream activators is essential in order to develop new strategies to target allosteric sites, either in p38 or its activators. The structures of individual MAPKs (*10–12*) and MAP2Ks (*13–16*) have been extensively studied and subsequent work has defined the interacting regions between these partners (*17–20*). The MAP2Ks contain a kinase interaction motif (KIM) at the start of their intrinsically disordered N-termini, which interacts with an allosteric docking site on their target MAPK. The KIM partially ensures specificity between members of the pathway (*17*), acts allosterically to expose the A-loop, preparing the MAPK for activation (*21, 22*), and also enhances the local concentration of kinase and substrate (*23*). However, there is little structural data on the global interactions between MAP kinase cascade components. Beyond the KIM interaction, molecular details of selectivity and activation of a MAPK by its upstream MAP2K remain unknown. Moreover, our knowledge of kinase-kinase interactions, in general, is restricted to homodimers and inactive conformations (*24–29*).

In order to further our understanding of how signals are transmitted through the MAP kinase pathway, we applied a multi-disciplinary approach, combining cryogenic electron microscopy (cryo-EM) with hydrogen-deuterium exchange mass spectrometry (HDX-MS), small angle X-ray scattering (SAXS), enhanced sampling molecular dynamics (MD) simulations and cellular assays, to characterise a complex between the MAP kinase p38α (MAPK14) and its activating MAP2K MKK6 (MAP2K6). Here, we present the first detailed molecular model of this highly dynamic and transient interaction between signalling components, providing insights into specificity and multi-step catalysis. Our findings open new routes to drug development for the important MAPK signalling pathway and also lead to a better understanding of a crucial step in kinase signalling cascades.

### Engineering an active and stable MKK6-p38**α** complex for structural studies

The MAP kinases are malleable proteins that have to adapt to multiple upstream and downstream effectors, as well as phosphorylate a large range of substrates. The proteins must therefore adopt multiple conformations and, crucially, interactions with effectors and substrates must be transient in order to maintain signal transmission. To stabilise the MKK6-p38α complex for structural studies, we created a chimera of MKK6, named MKK6^DD^GRA, where we replaced the native KIM with that from the *Toxoplasma gondii* effector protein GRA24 (*30*) that has a 100-fold higher affinity for p38α, while inducing the same conformation as the MKK6 KIM (Fig. S1A) (*31*). We also inserted phosphomimetic mutations in the MKK6 A-loop (S207D and T211D) to transform the kinase into a constitutively active form, ready to phosphorylate p38α (Fig. 1A and S1B and C). Analysis by SAXS showed significant heterogeneity in the size of the complex formed with p38α (Fig. S2A). By screening MKK6^DD^GRA with a combination of different nucleotide analogues and p38α A-loop mutants, we identified the mutant p38α^T180V^ (one of the phosphosites of the A-loop) in combination with the transition state analogue ADP.AlF ^−^, as the complex with the smallest radius of gyration (Rg) (Table S1). As a smaller Rg is associated with less flexibility, we reasoned that in this sample, the transition state analogue complex is stably assembled with the single residue available in the A-loop (Y182) and should be suitable for structural studies. Subsequent analysis, including HDX-MS, MD and Bayesian modelling confirmed that the engineering increases the population of the transphosphorylation-competent state of the complex and that its nature is consistent with that of the wild-type (see below).

**Fig. 1.**
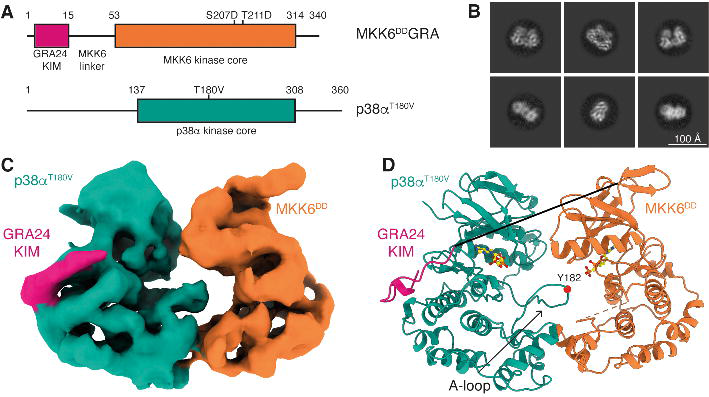
Structure of the MKK6^DD^GRA-p38 ^T180V^ complex. (**A**) Schematic of the two protein constructs. Key domains and mutations are indicated. (**B**) Representative 2D class averages from single-particle cryo-EM analysis of the MKK6^DD^GRA-p38α^T180V^ complex. (**C**) Segmented single-particle reconstruction cryo-EM map of the MKK6^DD^GRA-p38α^T180V^complex, resolved to 4 Å resolution and coloured according to (A). (**D**) Model of the MKK6^DD^GRA-p38α^T180V^ complex showing the overall structure of the complex. MKK6^DD^GRA-p38α^T180V^is represented as a cartoon and coloured according to (A), AMP-CP nucleotides are represented as balls and sticks (carbon atoms yellow). The MKK6 linker and A-loop are disordered and respectively shown as a full black line and an orange dashed line. The position of the Y182 residue in the p38α A-loop is indicated by a red sphere.

### Architecture of the MKK6-p38**α** complex

We imaged the 80 kDa MKK6^DD^GRA-p38α^T180V^complex by cryo-EM (Fig. 1). The purified complex was well dispersed and yielded 2D class averages that showed clear secondary structure features (Fig. 1B). The selected particles allowed the 3D reconstruction to a nominal resolution of 4 Å (map range 2.7-6.5 Å, Fig. 1C and S3, Movie S1). The final resolution was limited not only by the small size of the complex, but also by the highly dynamic nature of the interaction between the kinases, leading to particle heterogeneity. Therefore, very strict selection of particles was necessary to capture this conformation.

To build the model of the complex, we docked structures of p38α and MKK6 into the map. As all structures of inactive p38α available have an unstructured or mobile A-loop, we determined the crystal structures of human p38α, and an inactive mutant (p38α^K53R^), in which the A-loops are well ordered, in the occluded conformation, in order to have a starting point for refinement of the extended conformation (Table S2). The available MKK6 structures are not in the active conformation (Fig. S4). We therefore modelled the structure from the active conformation of MKK7 (*16*). Docking of these structures was followed by one round of morphing and real space refinement. We then computed the cross-correlation between experimental and predicted maps for relevant structures obtained from the ensemble obtained by combining Bayesian inference with MD simulations and SAXS data to select the model that best recapitulates the available data (Fig. 1D and S3C, Tables S3 and S4). The reconstruction shows the kinases adopting a ‘face-to-face’ conformation with most contact between the C-lobes. MKK6 is in the active kinase conformation based on its αC-helix rotated position and the DFG-in BLA-conformation according to kinase nomenclature (*32*) (Fig. S4), and density can be observed for nucleotide in its active site (Fig. S5). The p38α A-loop is ordered and extends toward the MKK6 active site; however, it remains dynamic, limiting the resolution of this region (Fig. S5). The p38α active site also binds nucleotide (Fig. S5), implying it is already in an active conformation. MKK6 interacts with the p38α CD site through the GRA24 KIM at the start of its N-terminal extension, which can be traced in the density (Fig. 1C and S5). Most of the linker between the KIM and the MKK6 kinase core, as well as the MKK6 A-loop, remains disordered and, therefore, unresolved. Interestingly, AlphaFold2 multimer (*33*) predicts a similar face-to-face conformation for the MKK6-p38α^WT^ complex, as well as for MKK6^DD^-p38α^WT^ and MKK6^DD^GRA-p38α^T180V^ (Fig. S6). Superimposing the predicted models to our experimental model shows a high consensus of the interaction. However, relative domain rotation in the prediction leads to differences in the interface of the two kinases (Fig. S6A). Remarkably, high confidence in intermolecular contact placement seen in the predicted aligned error (PAE) plots (Fig. S6D to F) support the overall face-to-face conformation of the complex.

The main kinase core fold interaction is between the αG-helix (residues 262-273) of MKK6 and a hydrophobic pocket in p38α, partly formed by the MAPK-specific insert in the C-lobe (Fig. 2A, C and D, Movie S2), that clamps the MKK6 αG-helix upon binding (Movie S3). This insert has been described as a lipid-binding site important in regulation (*34, 35*) and several studies have identified small molecules that target this pocket (*36–40*) (Fig. S7). Recently, it has also been shown to be important in MAPK substrate binding (*41, 42*) (Fig. S7). The exposed residues of this helix, as well as residues lining the MAPK hydrophobic pocket, are highly conserved in the MAP2Ks and MAPKs, respectively, suggesting a common interaction site for the pathways (Fig. 2E). Additionally, there is a potential interaction between the N-lobes via the loop between the β3-sheet and αC-helix of MKK6 (residues 87-89) and β-strands 1, 2 and 3 of the N-lobe of p38α (Fig. 2B). We validated the observed interactions using HDX-MS (*43*). In both the MKK6^DD^ (with the wild type KIM) and chimera MKK6^DD^GRA, several regions were protected from hydrogen/deuterium exchange in the presence of an excess of p38α: the N-terminus, the β3-strand, the A-loop and the αG-helix (Fig. 2G and S8 and Table S5). The interaction sites identified in solution highly correlate with the cryo-EM model for both the wild type and chimeric complexes. While no peptide was detected for MKK6 KIM, the GRA24 KIM was strongly protected upon binding with p38α. In an excess of MKK6^DD^ only the p38α MAPK insert and a small region of the N-lobe are protected from exchange (Fig. 2G and S8). We also evaluated the protection on p38α when we mutated the MKK6 αG-helix residues facing the p38α hydrophobic pocket to alanine (F264A, Q265A, L267A, K268A, E272A – referred to as MKK6^DD^ αG-helix mutant). Unlike with MKK6^DD^, the p38α MAPK insert is not protected in the presence of the MKK6^DD^ αG-helix mutant, which implies that this mutant cannot interact with p38α in that region (Fig. S9). Free energy calculations of p38α-MKK6 and p38α-MKK6 αG-helix mutant further support the lower stability of the C-lobe interface in the latter (“Additional text” in the Supporting Information and Fig. S9). These experiments define the αG-helix – MAPK insert interaction as the primary site of interaction between the two kinases during phosphorylation of the p38α A-loop.

**Fig. 2.**
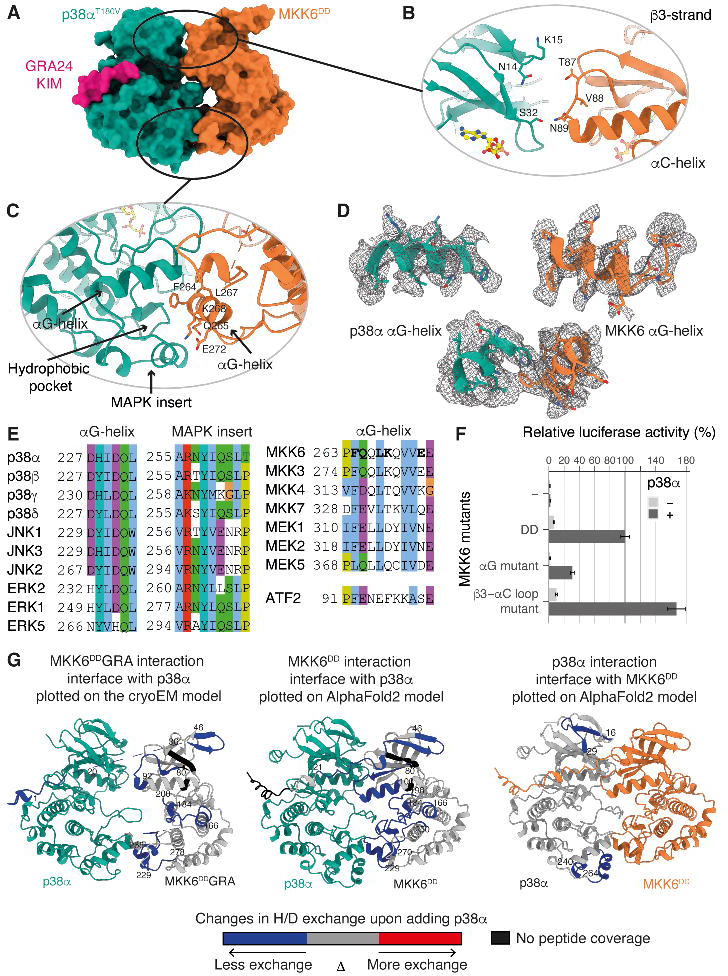
Novel interaction interfaces between MKK6 and p38α distal from the active site. (**A**) Surface representation of the MKK6^DD^GRA-p38α^T180V^ complex. (**B**) Potential interaction between the N-lobes. (**C**) Interaction between the αG-helix of MKK6 and the hydrophobic pocket of p38α. (**D**) αG-helices of p38α and MKK6 in the sharpened Coulomb potential map (black mesh). (**E**) Sequence alignment of MAPK hydrophobic pockets, MAP2K αG-helices and the p38α substrate ATF2 peptide (Clustal colouring scheme). (**F**) Luciferase reporter assay to monitor the activity of the p38α signalling pathway in HEK293T cells, showing the ability of MKK6^DD^ mutants to activate p38α (statistical analysis in Table S6). (**G**) MKK6^DD^GRA (left) and MKK6^DD^ (middle) interaction sites with p38α, and p38α interaction sites with MKK6^DD^(right), identified by HDX-MS. Regions showing protection upon the addition of p38α or MKK6^DD^, indicative of interaction, are highlighted in blue.

To further study the importance of these interactions on p38α activation, we used the MKK6^DD^ αG-helix mutant and also mutated the β3-αC loop of the MKK6 N-terminal interaction to alanine residues (T87A, V88A, N89A – referred to as MKK6^DD^ β3-αC loop mutant) and read out the effect on p38α signalling *in cellulo* using a reporter luciferase assay (Fig. 2F). We transiently transfected HEK293T cells with p38α and MKK6^DD^ mutants, together with a plasmid encoding firefly luciferase under the control of the AP-1 promoter, where AP-1 activity is stimulated by p38α via several substrates (e.g. ATF2, see Materials and Methods). The MKK6^DD^ αG-helix mutant reduced p38α signalling by 70% when compared to MKK6^DD^, demonstrating the important role this interaction plays in p38α activation. The MKK6^DD^ β3-αC loop mutant increased signalling to ∼170% over MKK6^DD^. This loop is the site of numerous activating mutations, particularly in cancer, that stabilise the ‘αC-in’ conformation (*44, 45*). This could explain the observed increase in signalling but does not rule out a role of the p38α N-lobe in the stabilisation of the active conformation of MKK6 in the face-to-face complex.

### Molecular dynamics confirm the complex is metastable and show how catalytically competent states can be formed

To better understand the details of the heterogeneous structural ensemble of the complex, and the mechanism of transphosphorylation of p38α on T180 and Y182 by MKK6, we ran a series of MD simulations, starting with a set of 18 simulations, each 1 μs long (Table S7). The simulations started from models derived from the cryo-EM structure: MKK6GRA-p38α (with WT A-loop) and MKK6-p38α, where we reverted the GRA24 KIM back to the WT sequence.

The simulations explored several conformations. In most cases, the contact between the KIM and p38α was retained, and catalytically competent states, where p38α Τ180 or Υ182 approached the γ-phosphate of MKK6 ATP at a catalytically compatible distance, were observed. Partial detachment of p38α from MKK6 was also observed in some instances. Additionally, the C-lobe interface proved to be important for the stability of the dimer, in agreement with our other observations. In all simulations in which the two kinases break apart, this tight hydrogen-bonding network at the C-lobe level is the last point of contact to be lost. In particular, the interaction between MKK6 αG-helix residues K268 and E272 and p38α S261 seems to be one of the main interactions keeping the two kinases in close proximity (Fig. S9E). This conformational variety is consistent with the SAXS and cryo-EM data, which are indicative of the transient nature of the interaction of the two kinases. With respect to the phosphorylation mechanism, in a number of trajectories, either p38α T180 or Y182 approach the ATP γ-phosphate located in the MKK6 active site (*46*) (Fig. 3 and Table S8, Movies S4 and S5). In particular, in the simulations of MKK6-p38α, both T180 and Y182 come close to the catalytic site. The tendency of monomeric, unphosphorylated p38α to adopt inactive, sequestered, A-loop conformations has been shown in previous-studies (*47*). Thus, the observation that the unphosphorylated A-loop of p38α adopts an exposed conformation in multiple instances is indicative of an active role of MKK6 in stabilizing such conformations. The active role of MKK6 is further confirmed by the fact that in the simulations in which the MKK6 KIM detaches, leading to the separation of the two kinases, the p38α A-loop transitions from an exposed to a sequestered conformation (Fig. S10).

**Fig. 3.**
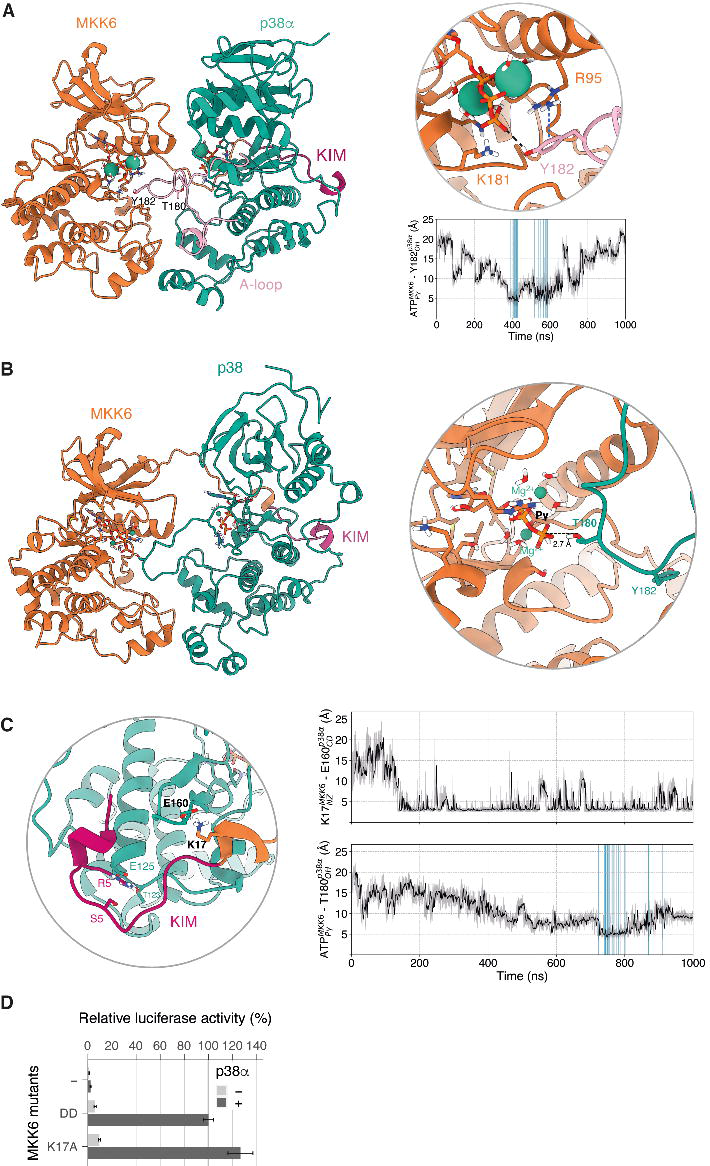
MD simulations show that both p38α Y182 and T180 can approach MKK6 ATP and that a rotated conformation of p38α favours T180 phosphorylation. (**A**) Frame extracted from one of the unrestrained MD simulations in which Y182 approaches the γ-phosphate of MKK6 ATP at a catalytically compatible distance (3.8 Å). The π-cation interaction of R95 with Y182 that further stabilizes Y182 close to ATP is shown in a blue dashed line. The frames where p38α Y182 reaches MKK6 ATP at a catalytically compatible distance in this set of simulations are highlighted in the plot of distances over time (blue). (**B**) Simulation frame in which p38α has rotated around its axis with respect to the cryo-EM structure. (**C**) Detailed view around the KIM. The frames where T180 reaches MKK6 ATP at a catalytically compatible distance in this set of simulations are highlighted in the plot of distances over time (blue). (**D**) Luciferase reporter assay to monitor the activity of the p38α signalling pathway in HEK293T cells showing the ability of MKK6^DD^ mutants to activate p38α (statistical analysis in Table S9).

### A rotated complex might facilitate the phosphorylation of p38**α** T180

In this initial set of MKK6GRA-p38α simulations, we observe some complexes in which T180 approaches the γ-phosphate of ATP in the catalytic site of MKK6 from an unexpected angle, where the p38α N-lobe underwent a 50-80° rotation around its axis (Fig. 3B, Movie S4). In this conformation, p38α was kept close to MKK6 through the KIM, despite the rotation. Before T180 gets close to the ATP, a key lysine located just after the KIM, K17, forms a salt-bridge with p38α E160 which seems to initiate the rotation of p38α (Fig. 3C).

To validate the functional relevance of this rotated intermediate observed in the simulations, we carried out site-directed mutagenesis, where we mutated the key lysine K17 to alanine. Interestingly, the MKK6^DD-K17A^ leads to a 26% increase in activity as measured by downstream signalling (Fig. 3D). The mutation is expected to abolish the K17-E160 salt bridge that primes the complex towards the rotated orientation (and T180 phosphorylation). As discussed below, we expect that by suppressing the rotated complex, the mutant could favour the formation of face-to-face dimers and enhance the more efficient phosphorylation pathway of Y182.

### Integration of simulations with SAXS data using Bayesian/Maximum entropy reweighting shows how the kinases associate

To better characterise intermediate states involved in the formation of the catalytically competent dimer, we extended the conformational sampling of the MD simulations using adaptive simulations (*48, 49*) (details in SI) and used a kinetic-based clustering of the total accumulated MD trajectories to determine the main metastable states. By extending the sampling to more than 18 µs and performing the kinetic clustering, we obtain five macro-states for p38α with MKK6GRA (Fig. 4 and Movies S6-S10). In the case of MKK6GRA-p38α, one of the clusters corresponds to an ensemble which contains conformations where the two kinases are face-to-face, equivalent to the cryo-EM structure (state A, Movie S6). Interestingly, in this macro-state, we also see conformations corresponding to the rotated N-lobe of p38α, in which T180 is ideally positioned for phosphorylation. The fact that the face-to-face and rotated conformations cluster within the same metastable state reflects the rapid interconversion from one to the other (few µs). In the second macro-state, the KIM is tightly bound, while MKK6 adopts different orientations (state E, Movie S7). The third macro-state represents conformations resulting from the detachment and non-specific binding of the two kinases, in which MKK6 binds to different surfaces of p38α (state C, Movie S8). Finally, we observe two macro-states in which the two kinases are held together through the C-lobe interface and in which the KIM is either bound (state B, Movie S9) or explores the N-lobe of p38α (state D, Movie S10). The connections between the macro-states (Fig. 4) indicate one or more reactive trajectories connecting one state to another. The resulting network shows that the binding of the KIM to its recognition site plays a fundamental role in establishing the specific contacts needed for the formation of the face-to-face dimer.

**Fig. 4.**
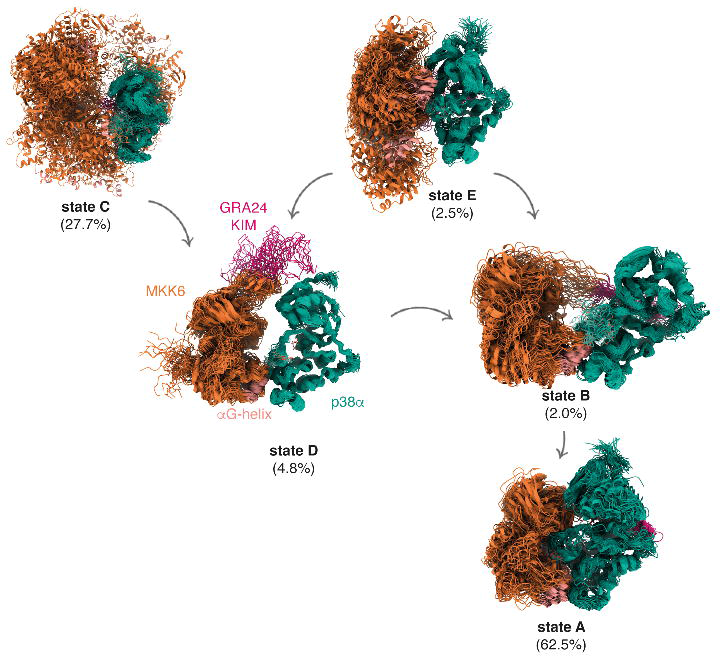
Kinetic-based clustering of the accumulated simulations of MKK6GRA-p38α. Each macro-state shows an overlay of 50 representative conformations sampled proportionally to the equilibrium probability of each micro-state in the corresponding macro-state corresponding to different kinetically distinct states. The population of each state as derived from the fitting to the MKK6^DD^GRA + p38α^WT^ + ADP + AlF ^−^ SAXS curve (Fig. S2) is given in parenthesis.

Despite a total sampling time of more than 18 µs, we could not fully converge the kinetics and the population of all the macro-states directly from the simulations, probably due to the slow nature of some of the transitions seen in protein-protein associations (*49*). An exhaustive sampling of complex formation from the detached states would require a prohibitive number of long MD simulations. We therefore refined the structural ensemble and validated the clustering results by using the SAXS data and a Bayesian/maximum entropy approach (*50–52*). The SAXS profile of each conformer within the clustered MD ensemble was calculated, and the weights associated with each conformer in the ensemble that maximise the agreement with experiments were iteratively determined (Fig. S11A). For the refinement of the MKK6GRA-p38α ensemble, we used the SAXS curves of the complex with ADP.AlF ^−^ and with the non-hydrolysable ATP analogue AMP-PCP. The reweighted ensemble fits extremely well with the SAXS curve of MKK6^DD^GRA-p38α^T180V^ with ADP.AlF ^−^ (χ^2^=0.89, Movie S15), which showed the most compact conformation (Table S1). Reassuringly, all the macro-states from kinetic modelling contribute to it (Fig. 4). State A, which contains conformations equivalent to the cryo-EM structure, is the most populated (62.5%), followed by state C (27.8%), which reflects the transient nature of the complex and the non-specific binding associated with such transient complexes, and finally ensemble E (2.5%). From the weights (and the other experiments), we deduce that the latter is the main intermediate for the formation of the face-to-face complex. After the KIM domain is bound, the MD simulations indicate a significant role of the hydrophobic patch and a network of hydrogen bonds located at the interface of the C-lobes of the two kinases in the formation of the complex. This is in accordance with the 70% decrease in activity seen in the MKK6^DD^ αG-helix mutant (Fig. 2F), as well as the metadynamics free energy calculations and the HDX-MS data (Fig. S9).

The ensemble reweighted to match the SAXS data obtained with the MKK6^DD^GRA-p38α^WT^with the ATP analogue AMP-PCP also resulted in a good fit (χ^2^=2.1). The population of the face-to-face cluster A decreases to 1% while that of the detached cluster C and intermediate D increases considerably, to 71.5% and 21.7%, respectively (Fig. S11B, C). While the populations of the macro-states shift, as expected as the transition state is not stabilised, the general description of the macro-states remains similar. Within cluster A, the weights of the ‘rotated’ face-to-face dimer increase (Fig. S11B). We also sampled the conformational space of the MKK6-p38α with the same adaptive MSM approach. The four macro-states found in the MKK6-p38α complex are equivalent to four of the macro-states found in MKK6GRA-p38α (Fig. S12), reflecting the similarity of the dynamics of the two complexes (see Supplementary text).

### The N-terminal extension of MKK6 remains mainly disordered and its length and secondary structure elements contribute to pathway specificity

Our structure and macro-state modelling imply that once the KIM is bound the N-terminal linker plays an important role in conformational sampling, allowing the catalytically competent state to form, a phenomenon that has been observed in other tethering proteins (*53*). There is a wealth of data on interactions between KIMs and MAPKs, but the role of the remainder of the N-terminus is poorly understood (*54, 55*). In our cryo-EM map, the engineered KIM is clearly visible (Fig. 1C and S5), but it does not appear to interact further with p38α. To investigate whether the N-terminal extension has a role beyond simply associating the kinase to the KIM, we again used the luciferase reporter assay. We produced constructs that scan the region between the KIM and the first consensus β-sheet of the kinase core with alanine blocks in order to determine if there is sequence specificity or direct interaction with p38α (Fig. 5). Our results show that the sequence of the middle region (MKK6^DD^ Ala scan 28-39) of the N-terminal linker has no effect on p38α signalling *in cellulo*. However, the sequence of the linker close to the KIM (MKK6^DD^ Ala scan 18-29) and the region close to the kinase core, which comprises the predicted β-strands (MKK6^DD^ Ala scan 38-49, Fig. 5) seem to have some importance as their mutation to alanines reduced p38α signalling by 27% and 58% respectively.

**Fig. 5.**
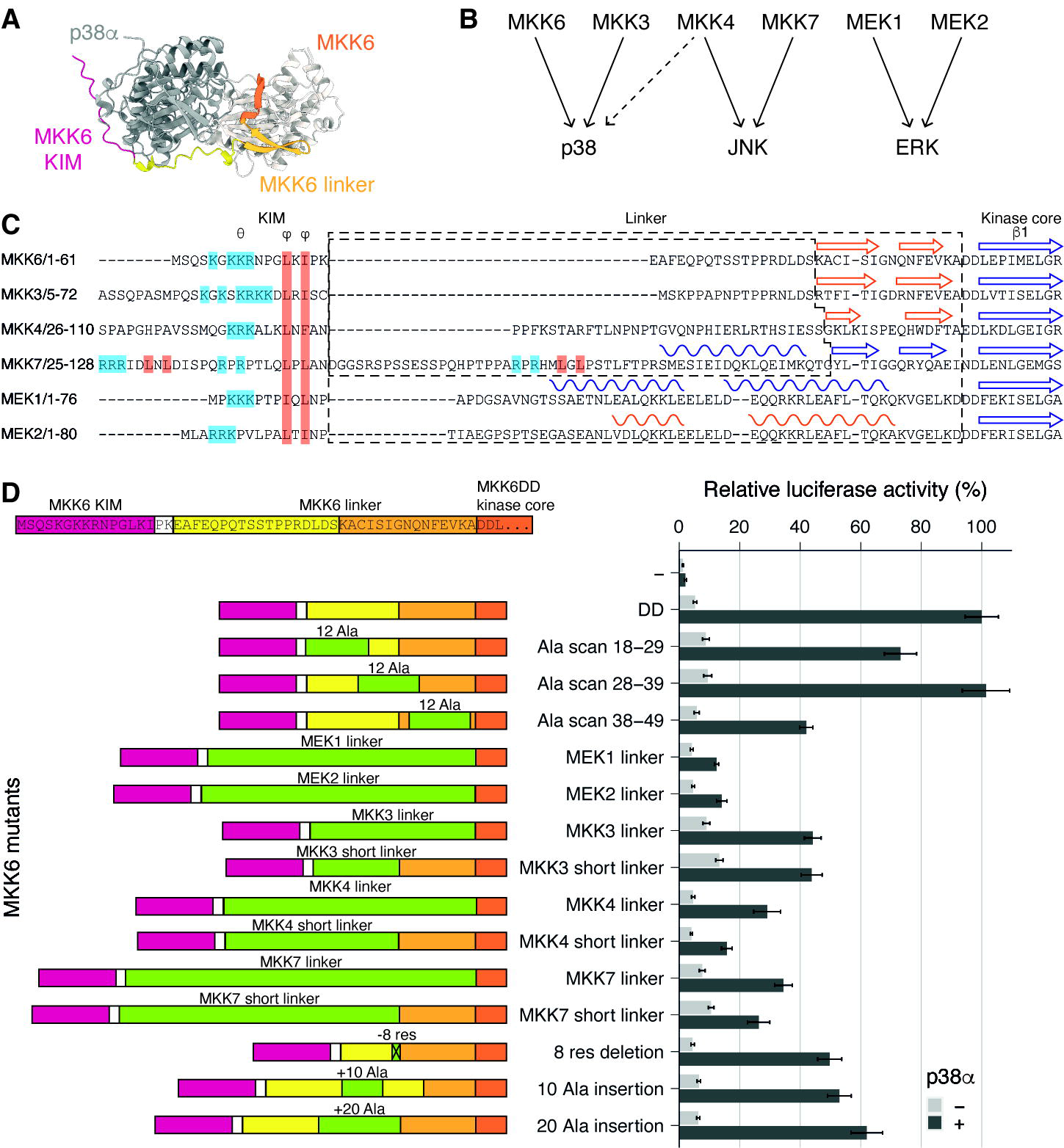
MKK6 N-terminal extension length and secondary structure define the specificity of p38α activation. (**A**) Top view of an MKK6-p38α AlphaFold2 multimer model, for illustrative purposes. (**B**) Specificity of the MAP2K/MAPK signalling pathways. (**C)** Sequence alignment of MAP2K N-termini. Secondary structure elements are indicated, from experimental structures in blue and AlphaFold2 predictions (pLDDT score > 65) in orange. (**D**) Luciferase reporter assay to monitor the activity of the p38α signalling pathway in HEK293T cells showing the ability of MKK6^DD^ mutants and chimeras to activate p38α (statistical analysis in Table S10).

While KIM sequences are important, they do not sufficiently explain specificity. Comparing the N-terminal linkers of the MAP2Ks, they differ significantly in their length and secondary structure elements (*16, 56, 57*) (Fig. 5B and C), could this contribute to specificity between the pathways? When we substituted the MKK6 linker with those from the MAP2Ks of the other pathways, activity was significantly reduced (Fig. 5D). This is most drastically observed with the linkers from MEK1 and 2. Even the linkers from MAP2Ks that activate p38α see reduced activity and are not sufficient to rescue the activation, even though the KIM, MAP2K and MAPK are in the correct combination. Our structure and macro-state models imply that the length of the linker is just sufficient to allow the observed interactions between the kinases, in particular, the MAP2K αG-helix position relative to the MAPK hydrophobic pocket formed by the MAPK insert. By removing or adding 10 residues to the linker region, activity is reduced by ∼50%. This implies that the length of the linker, controlled by either the number of amino acids and/or secondary structure elements, is important in the positioning of the MAP2K for engagement with the MAPK and is finely tuned for MAP kinase pairs. The specificity of a MAP2K is therefore defined by cooperation between the KIM, the linker and the kinase core folds themselves.

## Discussion

### Initial docking

The N-termini of the MAP2Ks contain a conserved KIM motif that is required for binding to their substrate MAPKs. The first step in activation is KIM binding after which a wide variety of conformations are observed, showing the fast timescale of association/dissociation. During the MD simulations, KIM binding was seen to induce allosteric changes in p38α, which adopts a prone-to-be-phosphorylated conformation with its A-loop extended, exposing the tyrosine and threonine residues, in agreement with crystal structures. Our data indicate that the interaction between the MKK6 N-terminus and p38α extends beyond the hydrophobic residues of the KIM and is important in the activation process. Although most of the MKK6 N-terminal linker remains disordered, its length seems to be tightly linked to the substrate MAPK and the presence of some secondary structural elements differs between MAP2Ks contributing to specificity (Fig. 5C). Our reweighted ensemble, obtained by combining Bayesian inference with adaptive MSM simulations and SAXS data, shows that once the KIM is bound, a large number of conformations are sampled before the engagement of the αG-helix (Fig. 4). This is fully consistent with the experiments that show that if the linker is not the correct length, the engagement of the αG-helix will be impeded. Once p38α is recruited, some parts of the MKK6 N-terminal linker could participate in the conformational shift towards the MKK6 active conformation, but its main function appears to be to prepare the MAPK A-loop and guide the engagement with the αG-helix.

### Engagement with substrate

We show that the predominant interaction during activation is mediated by the αG-helix of the MAP2K MKK6 and the p38α hydrophobic pocket formed by the αG-helix of p38α and the specific MAPK insert. This region is also important in MAPKs for substrate recognition (*41*) and interaction with scaffold proteins and downstream regulators such as phosphatases (*27, 42, 58*). The engagement of αG-helices seems to be an emerging theme in kinase heterodimers: only a few structures have been determined, such as the KSR2-MEK1 heterodimer (*27*) and more recently, the BRAF-MEK1 complex (*25*), both upstream components of the related ERK MAPK pathway. In both these structures the αG-helices interaction is very similar, as well as the relative orientation of the kinases. Several studies have shown that small molecules can target this region in p38 isoforms (*36–40*) and modulate the position of the MAPK insert helices (Fig. S7). However, as the pocket is very deep, the molecules may not extend far enough to disrupt the interactions identified here. There has been intense interest, and some success (*59*), in developing drugs that target p38α, mostly binding to the nucleotide pocket, but they have been mired by off-target and toxicity effects (*9*). The molecules that have been identified as binding to the MAPK insert pocket could be further developed to better disrupt both the MAP2K interaction, and that of substrates, which could lead to highly specific p38 inhibitors.

The MKK6 KIM–p38α docking site, the MKK6 N-terminal linker, and the MKK6 αG-helix–p38α hydrophobic pocket interactions, are clearly essential for defining the specificity of the kinase-substrate interaction, for positioning the two kinases, and for triggering the necessary changes towards the MKK6 active kinase conformation and positioning the A-loop of p38α to be accessible for phosphorylation. Our data demonstrate that altering any of these three components perturbs the activation of downstream signalling.

### Freedom of interaction at the catalytic centre

All contacts observed between the kinases are distal to the active site, explaining the lack of sequence specificity or conservation in MAPK A-loops in the region preceding the TxY motif (*57*) and our HDX-MS data show no tight interaction between the A-loop of p38α and MKK6 (Fig. 2 and S8). Rather than the active site of a classical enzyme, where substrates are perfectly positioned for catalysis, the MAP2K-MAPK complex appears to provide a zone of proximity, allowing the flexible activation loop to move, but increasing the probability, or local concentration, of the p38α T180 and Y182 residues to be positioned for nucleophilic attack of the γ-phosphate of ATP by relative positioning of the two kinases. Dual specificity MAP2Ks are unusual as the two amino acids targeted are significantly different to each other, when compared to serine/threonine kinases. In addition, the two substrate phosphorylation sites are only one residue apart and the MAP2Ks need to have an active site that can accommodate, effectively, four substrates (A-loop **T**-x-Y; T-x-**Y**; **T**-x-Y^P^; T^P^-x-**Y** (targeted residue in **bold**)), all with significantly different sizes and charges. This implies that flexibility in the active site is essential in order to accommodate such a wide variety of substrates. By not binding any residues of the A-loop specifically, the various different residues and phosphorylation states could be accommodated by sacrificing catalytic efficiency for flexibility.

The MD simulations revealed that the face-to-face architecture of the hetero-kinase dimer of our model is compatible with the phosphorylation of either T180 or Y182 of the p38α A-loop (Fig. 3 and S13). Another conformation, where p38α rotated, emerged and seems to favour the phosphorylation of T180. Kinetic data have shown that the dual phosphorylation of p38α by MKK6 involves a partially processive mechanism *in vitro,* in which the monophosphorylated intermediates can either dissociate from the enzyme or proceed to the next catalytic step. Wang *et al. (60)* and others (*61*) propose that both p38α monophosphorylated forms can be produced by MKK6 catalysis with a preference for Y182, whose phosphorylation is four-fold faster than that of T180. With respect to the second catalytic step, the experimentally measured kinetic rates indicate that phosphorylation at Y182 in the first step enhances the catalytic efficiency of MKK6 phosphorylation at T180 in the second step. According to our simulations (Fig. 3 and S13), when the N-lobe of p38α undergoes a large conformational change towards the rotated dimer, T180 is in the right position to be phosphorylated, while Y182 is more distant from the catalytic site, which might explain the observed lower catalytic efficiency of MKK6 when the second phosphorylation is on Y182 by kinetic measurements. Conversely, Y182 seems to be able to easily approach MKK6 ATP in the face-to-face catalytically-competent conformations and then be phosphorylated. Thus, the difference in the experimental kinetic rates measured for the first phosphorylation step may reflect the rotation of the p38α N-lobe around its axis, which seems to be involved in the T180 phosphorylation only. The partial processivity of the dual phosphorylation in a cellular context remains an open question (*62, 63*), but the architecture of the interaction we observe between MKK6 and p38α, as well as the micro-states seen in modelling seems to be compatible with either a distributive, partially processive or fully processive mechanism.

By combining cryo-EM and MD simulations, together with HDX-MS and structure-driven mutagenesis *in cellulo*, we describe the range of attributes that lead to MKK6 selectivity and p38α activation. Moreover, we identify critical interfaces for dimer stabilisation. In this regard, small molecules that prevent p38α-MKK6 complex dimerisation could be therapeutically effective in treating diseases resulting from abnormal p38α-ΜΚΚ6-mediated signalling. What is more, our data describe the first molecular details of the activation of one protein kinase by another – one of the most fundamental mechanisms in cell signalling.

## Materials and Methods

### Plasmids

Plasmids were ordered from GenScript (gene synthesis, cloning and mutagenesis). For recombinant protein expression, human p38α sequences (WT, T180V, Y182F, and K53A mutants) were fused to a His6 tag with a 3C protease cleavage site and cloned into a pET-28b vector; MKK6^DD^GRA sequence (constitutively active S207D T211D mutant of human MKK6 with GRA24 KIM) was fused to a twin StrepII tag with a 3C cleavage site and cloned into a pFastBac1 vector. For cellular reporter assays, p38α and MKK6^DD^ mutants sequences were cloned into pcDNA3.1 plasmids. The lambda Phosphatase plasmid was obtained from Addgene (a gift from John Chodera & Nicholas Levinson & Markus Seeliger; Addgene plasmid #79748; http://n2t.net/addgene:79748 ; RRID:Addgene_79748) (*64*).

### Protein expression and purification

Constructs of p38α were co-transformed with the lambda phosphatase plasmid into Rosetta™(DE3)pLysS *E. coli* competent cells (Novagen) with appropriate antibiotics. Cells were grown in LB at 37°C until the OD600 = 0.6-0.8, induced with 0.5 mM IPTG, incubated at 16°C overnight, and harvested by centrifugation.

The MKK6^DD^GRA constructs were transformed into DH10 EMBacY *E. coli* cells to produce recombinant baculoviruses subsequently used for protein expression in *Sf21* insect cells (*65*). Cells were harvested 48h after proliferation arrest by centrifugation.

Cell pellets were resuspended in lysis buffer (50 mM HEPES pH 7.5, 200 mM NaCl, 10 mM MgCl_2_, 5% glycerol, 0.5 mM TCEP, with a Pierce protease inhibitor EDTA-free tablet (Thermo Scientific) and a trace of DNaseI (Sigma)). Cells were lysed by sonication on ice and the lysate cleared by centrifugation.

For p38α, supernatant was loaded onto a pre-packed 5 ml HisTrap column (GE Healthcare), equilibrated according to the supplier’s protocols with wash buffer (50 mM HEPES pH 7.5, 200 mM NaCl, 10 mM MgCl_2_, 5 % glycerol, 0.5 mM TCEP) with 1% elution buffer (wash buffer with 500 mM Imidazole). Tagged-protein was eluted with elution buffer and p38α-containing fractions were pooled, incubated with 3C-protease and dialysed against wash buffer at 4°C overnight. The sample was run through the HisTrap column again to remove uncleaved protein and flow-through fractions were collected.

For MKK6^DD^GRA, supernatant was loaded onto a pre-packed 5 ml StrepTactin XT column (IBA), equilibrated according to the supplier’s protocols with wash buffer. Tagged-protein was eluted with elution buffer (wash buffer with 50 mM Biotin), and MKK6-containing fractions were pooled, incubated with 3C-protease and dialysed against wash buffer at 4 °C overnight. The sample was run through the StrepTactin XT column again to remove uncleaved protein, HisTrap column to remove the protease, and flow-through fractions were collected.

Pure proteins were concentrated using 10 kDa Amicon Ultra membrane concentrators (Millipore), flash-frozen in liquid nitrogen and stored at −80℃.

In order to prepare the hetero-kinase complex, MKK6^DD^GRA and p38α aliquots were thawed, mixed in 1:1 mass ratio and incubated on ice for 1h. The sample was then concentrated to 10 mg/ml with a 10 kDa Amicon Ultra membrane concentrators (Millipore) and loaded onto a Superdex S200 increase 10/300 GL size-exclusion column (GE Healthcare) equilibrated in wash buffer (with a reduced glycerol concentration of 2% when for cryo-EM studies). Purest peak fractions were pooled, flash-frozen in liquid nitrogen and stored at −80℃ before being used for structural studies or biochemical assays.

### SAXS

MKK6^DD^GRA-p38α complexes purified by gel filtration were complemented with either 10 mM ADP, 10 mM NH_4_F and 1 mM AlCl_3_ or 10 mM AMP-PCP and incubated on ice for 30 minutes before proceeding.

Small Angle X-ray Scattering data were collected at the bioSAXS beamline B21 at the Diamond Light Source (DLS), UK (proposal SM23091-1). Scattering curves were measured from solutions of the different MKK6^DD^GRA/p38α samples in SAXS buffer (50 mM HEPES pH 7.5, 200 mM NaCl, 10 mM MgCl2, 5 % glycerol, 0.5 mM TCEP). Buffer subtractions and all other subsequent analyses were performed with the program Scatter IV (*66*) and SAXS profile computation used FoXS (*67*) (Table S11).

### Cryo-EM specimen preparation and data collection

Purified MKK6^DD^GRA-p38α^T180V^ was complemented with 250 µM AMP-CP, 250 µM NH_4_F and 25 µM AlCl_3_, and incubated on ice for 30 minutes before proceeding. UltraAufoil 1.2/1.3 grids were glow-discharged for 30 seconds at 25 mA (PELCO easy glow). During the vitrification procedure on a Vitrobot Mark IV (FEI), 4 μl of sample at 6 μM concentration was applied to the grid. The blotting force was set to 0 for a total time of 3.5 seconds. Grids were then clipped and screened on a FEI Talos Glacios electron microscope (EMBL Grenoble) operating at 200 kV. Selected grids were then sent for data collection on a FEI Titan Krios (EMBL Heidelberg) operating at 300 kV. Micrographs were automatically collected using SerialEM (*68*) from a K2 Quantum detector (Gatan) and a GIF Quantum energy filter (Gatan), at a nominal magnification of 215 kx (corresponding to 0.638 Å/pixel at the specimen level). Three different data collections with a total of 28,633 movies of 40 frames were collected in electron counting mode with a defocus range of −1.5 to −3 µm, 0.1 µm step, with a total exposure of 62.77 e^−^/Å^2^.

### Cryo-EM processing

Each dataset was treated independently following a similar workflow and the best particles corresponding to the hetero-kinase sample were finally combined and further processed.

Movies were imported into cryoSPARC (*69*). All movie frames were aligned and motion-corrected with Patch Motion correction and CTF estimated with Patch CTF. Data were manually curated to eliminate micrographs with large motions, poor resolution, and high ice thickness. The resulting micrographs showed a low signal/noise ratio, but classical picking with blob picker failed to identify particles. We successfully picked a large number of particles by using a combination of Topaz particle picker (*70*) and template picker. Particles were extracted with a box size of 300 x 300 pixels and 2D classification was used to eliminate junk and the noisiest particles. We then applied *ab-initio* and heterogeneous refinement, requesting 3-6 models, and particles corresponding to the hetero kinase model were selected for further 2D classification. Between 2-3 rounds of 2D classification were necessary to select good particles, showing secondary structural features and low background noise. Non-uniform refinement was then used to obtain a 5 Å resolution map. An exemplary workflow is shown in Figure S3A for dataset 2.

To improve resolution, we then combined selected particles from each dataset (86,510 particles in total) and we applied 2D classification, *ab-initio*, heterogeneous and non-uniform refinement to obtain a final map at 4.00 Å resolution, corresponding to 35,123 particles (Fig. S3B). Maps were sharpened in cryoSPARC and using locscale (*71*).

The initial model was built from the combination of different structures: the crystal structure of p38α WT presenting an ordered activation loop (PDB-5ETC), the crystal structure of p38α bound to GRA24 KIM peptide (PDB-5ETA) (*31*), and the homology model of MKK6^DD^ based on the crystal structure of the active MKK7^DD^ structure (PDB-6YG1) (*16*) obtained with SWISS-MODEL (*72*). The model of each kinase was individually docked into the map as a rigid body using the phenix.place.model and then refined with one round of morphing and real space refinement using Phenix (*73*) and corrected in COOT (*74*). To validate the rebuilt model, we relied on cross-correlation calculations between the experimental maps and those predicted from the MD simulations to select the model that best recapitulates the available data. We considered a set of 87 structures including the rebuilt model, AF2 predictions, complex structures including the linker and p38α activation loop modelled using the *automodel* functionality of MODELLER, and 79 clusters extracted from one relevant MD simulation showing p38α Y182 close to MKK6 ATP (rep4 with restraints). Given the sensitivity of cross-correlation calculations to atomic B-factors, we re-assigned temperature factors for each structure using Phenix by running one round of refinement with default parameters. This step allowed us to compare the cross-correlation scores assigned to all the models in the dataset. Cross-correlation scores were then computed considering the entire complex and a selection of key structural features, i.e., hydrophobic patch, KIM, p38α activation loop, and the MKK6 region close to the resolved linker (details in Table S3). By selecting a specific region of the structure, cross-correlation was computed in the area surrounding the selection only. Considering the entire complex, the rebuilt model was assigned the highest cross-correlation score. However, one of the predicted models was assigned to a higher score than the rebuilt model for the area around the hydrophobic patch, the p38α activation loop, and the MKK6 region close to the linker. Therefore, we incorporated the structural information for these features into the final model as they are in better agreement with the experimental data. Table S3 summarizes the cross-correlation scores for all structures. See Table S4 for data processing and refinement statistics. Figures were prepared in ChimeraX (*75*) and validation performed with Phenix Comprehensive Validation.

### Crystallisation, data collection and structure determination of inactive p38L J with a stable A-loop

Crystallization conditions were established by testing several commercial screens at the EMBL High Throughput Crystallisation Laboratory. Crystals of p38α^K53R^were obtained at 20LJC by the sitting drop method from solutions containing 10 mg/ml p38α^K53R^in 20 mM Tris HCl pH 7.5, 150 mM NaCl, 10mM MgCl_2_, 1mM DTT, 5% (v/v) glycerol, equilibrated against 3.5 M Na Formate pH 7.0. Crystals were transferred to a cryoprotection buffer prepared by equilibrating protein buffer solution against reservoir solution supplemented with 20% (v/v) glycerol in a sitting drop plate and harvested using a micromesh loop (MiteGen, Ithica, NY, USA), plunged into liquid nitrogen and stored at 70 K. Crystals of wild type p38α were obtained as above but equilibrated against 1.5 to 1.6 M NH_4_(SO_4_)_2_, 50mM MgCl_2_ and 50 mM TRIS/HCl pH 8.0. Crystals were harvested directly from the mother liquor (*76*) using a micromesh loop (MiteGen, Ithica, NY, USA), plunged into liquid nitrogen and stored at 70 K. Diffraction data were collected at beamlines ID23-1 and ID23-2 at the ESRF (Grenoble, France) on an ADSC Q315 CCD detector (ID23-1) or a MAR225 CCD detector (ID23-2) to between 2.2 and 2.8 Å resolution. Crystals of p38α and p38α^K53R^ contained multiple leaves of crystals and the combination of a microfocussed or micro beam in combination with automated mesh scans was essential. Both p38α and p38α^K53R^ crystallised in the orthorhombic space group *P*2_1_2_1_2_1_ with one molecule in the asymmetric unit and similar unit cell dimensions (Table S2). Data were processed with XDS (*77*) and programs from the Collaborative Computational Project Number 4 suite (*78*). The structures were determined by molecular replacement using MolRep (*79*). For the p38α^K53R^ structure, accession code 1WFC (*10*) was used as search model with all water molecules removed. For p38α the p38α^K53R^ structure was used as a search model. Refinement was carried out alternately using Phenix (*80*) and by manual rebuilding with COOT (*74*). Models were validated using Molprobity (*81*).

### AlphaFold2 predictions

AlphaFold2 multimer (*33*) was run with a local ColabFold (*82*) set-up available from https://github.com/YoshitakaMo/localcolabfold. Input sequences of MKK6 + p38α complexes were provided in csv format and run with following option settings: --num-recycle 3 --model-type AlphaFold2-multimer-v2 --num-models 5. Resulting models and PAE plots were coloured using ChimeraX (*83*).

### HDX-MS

HDX-MS experiments were performed at the UniGe Protein Platform (University of Geneva, Switzerland) following a well-established protocol with minimal modifications (*84*). Details of reaction conditions and all data are presented in Table S5. HDX reactions were done in 50 LJl volumes with a final protein concentration of 5.4 μM of MKK6. Briefly, 272 picomoles of protein were pre-incubated with 815 picomoles of p38α or corresponding buffer in 5 µl final volume for 5 min at 22 °C before initiating the reaction.

Deuterium exchange reaction was initiated by adding 45 µl of D_2_O exchange buffer (20 mM Hepes pH 7.5 / 200 mM NaCl / 10 mM MgCl_2_ / 500 nM TCEP / 2 mM AlCl_3_ / 1 mM NH_4_F / 1 mM AMP-CP supplemented with 6 LJM of p38 or equivalent buffer) to the protein mixture. Reactions were carried-out at room temperature for 3 incubation times in triplicates (3 s, 30 s, 300 s) and terminated by the sequential addition of 20 μl of ice-cold quench buffer 1 (4 M Gdn-HCl / 1 M NaCl / 0.1 M NaH_2_PO_4_ pH 2.5 / 1 % Formic Acid (FA) / 200 mM TCEP in D_2_O). Samples were immediately frozen in liquid nitrogen and stored at −80 °C for up to two weeks.

To quantify deuterium uptake into the protein, samples were thawed and injected in a UPLC system immersed on ice with 0.1 % FA as liquid phase. The protein was digested via two immobilized pepsin columns (Thermo #23131), and peptides were collected onto a VanGuard precolumn trap (Waters). The trap was subsequently eluted, and peptides separated with a C18, 300Å, 1.7 μm particle size Fortis Bio 100 x 2.1 mm column over a gradient of 8 – 30 % buffer over 20 min at 150 LJl/min (Buffer B: 0.1 % formic acid; buffer C: 100 % acetonitrile). Mass spectra were acquired on an Orbitrap Velos Pro (Thermo), for ions from 400 to 2200 m/z using an electrospray ionization source operated at 300 °C, 5 kV of ion spray voltage. Peptides were identified by data-dependent acquisition of a non-deuterated sample after MS/MS and data were analyzed by Mascot. All peptides analysed are shown in Table S5. Deuterium incorporation levels were quantified using HD examiner software (Sierra Analytics), and quality of every peptide was checked manually. Results are presented as percentage of maximal deuteration compared to theoretical maximal deuteration. Changes in deuteration level between two states were considered significant if >15% and >1.5 Da and p< 0.01 (unpaired t-test). The mass spectrometry proteomics data have been deposited to the ProteomeXchange Consortium via the PRIDE (*85*) partner repository with the dataset identifier PXD040499 and 10.6019/PXD040499.

### AP-1 reporter assay

HEK293T cells were maintained in DMEM medium (DMEM) supplemented with 10% (v/v) FBS (Thermo Fisher), at 37 °C and 5% CO_2_. According to the manufacturer’s protocol, the AP-1 firefly luciferase reporter vector and the constitutively expressing *Renilla* luciferase vector (BPS Bioscience #60612) were transiently co-transfected together with MKK6^DD^ mutants and p38α pcDNA3.1 plasmids (or empty vector for controls) with Lipofectamin2000 (Thermo Scientific) (Fig. S14). Firefly and *Renilla* luciferase activities were measured 24h post-transfection using the Dual Luciferase assay system (BPS Bioscience #60683) with a Clariostar microplate reader (BMG Germany). Each experiment was repeated three times (biological replicate) and each time, three transfections per condition were performed (technical replicate), for a total of nine experiments per construct. Each firefly luciferase value was normalised by its corresponding *Renilla* luciferase measure to correct for cell quantity and transfection efficiency. The luciferase activity was expressed relative to the replicate positive control (MKK6^DD^ + p38α) intensity. Statistical analysis on the AP-1 reporter assay was performed using Rstudio. Briefly, normal distribution was tested with a Shapiro Wilck test and variance homogeneity with a Levene’s test. To compare the mutants to the control (MKK6^DD^ + p38α), either a Welch t-test or a Wilcoxon rank sum exact test was performed.

### Mass spectrometry

Purified MKK6^DD^GRA-p38α complexes were complemented with 10 mM ATP. Intact mass was measured on a quadrupole time of flight (Q-TOF) Premier mass spectrometer (Waters/Micromass) coupled to an Acquity UPLC System (Waters Corporation). Solvent A was water, 0.1% formic acid, and solvent B was acetonitrile, 0.1% formic acid. Samples were acidified using 1% TFA prior to injection onto an Acquity UPLC Protein BEH C4 column (Waters Corporation). Acquisition was carried out using the standard ESI source in positive ion mode over the mass range 500–3500 m/z. Data were externally calibrated against a reference standard of intact myoglobin, acquired immediately prior to sample data acquisition. Spectra from the chromatogram protein peak were then summed and intact mass was calculated using the MaxEnt1 maximum entropy algorithm (Waters/Micromass).

PTMs were measured by LC-MS/MS. SP3 protocol (*86*) was used for sample preparation. Analysis was performed on an UltiMate 3000 RSLC nano LC system (Dionex) fitted with a trapping cartridge (µ-Precolumn C18 PepMap 100) and an analytical column (nanoEase™ M/Z HSS T3 column 75 µm x 250 mm C18, 1.8 µm, 100 Å, Waters) coupled to a Fusion Lumos (Thermo) mass spectrometer using the proxeon nanoflow source in positive ion mode. Acquired data were processed by IsobarQuant (*87*), as search engine Mascot (v2.2.07) was used.

### Protein structure preparation for MD simulations

The model derived from cryo-EM of p38α in complex with the MKK6^DD^GRA was used as a starting conformation for the unbiased MD simulations. The bound ADP that was resolved in the cryo-EM structure was replaced by ATP, which was docked to the binding sites of p38α and MKK6 using the Glide software (*88*) (Schrödinger Release 2021-4). In particular, the docked ATP maintained the ADP pose seen in the cryo-EM structure, while the position of AlF_4_ ^−^ in the binding site of p38α was used to guide the position of the γ-phosphate group of the docked ATP. The same γ-phosphate position was used for the ATP in the binding site of MKK6, even though AlF_4_ ^−^ was not present in this binding site. The mutated residues S207D and T211D on the resolved MKK6^DD^ were reverted back to the WT ones. The K53R mutation was introduced unintentionally when building the model of p38α with the WT KIM, as this was the mutant used in the crystal structure. It is a kinase-dead mutation that stabilizes the protein for structural studies but has no influence on the studies presented here. The structure of the p38α-MKK6GRA produced by the aforementioned procedure was then used to model the N-terminal tail of the WT MKK6 (residues M1-K14 of the MKK6 WT sequence) using the crystal structure of p38α bound to the WT MKK6 KIM (*31*) and with the *automodel* functionality of MODELLER (*89*).

### MD simulations setup

Prior to any simulation, the protonation state of each residue at pH 7.4 was calculated using the PROPKA 3.1 algorithm as implemented in the PlayMolecule web application (*90*), which left all residues in their usual charge states, except for p38α H228, which was double protonated. For all simulated systems, DES-Amber force field was used for the protein as it has been shown to be able to capture the dynamics of protein-protein complexes more accurately than other force fields to this date (*90*). For the ATP coordinated Mg^2+^ ions, optimized Mg force field parameters that yield water exchange on timescales that agree with experiments (*90*) were combined with the protein force field. The Meagher et al. parameterization for ATP (*91*) was used, as these parameters have been benchmarked recently and shown to be able to reproduce the experimentally observed coordination modes of ATP (*92*). Each system was enclosed in an octahedral box with periodic boundary conditions and solvated with TIP4PD water molecules as described in the DES-Amber force field, while Na^+^ and Cl^−^ ions were added to reach neutrality at the final NaCl concentration of 150 mM.

The MD simulations were performed using the GROMACS 2021.3 simulation package (*93*) patched with the PLUMED 2.4.1 plug-in (*94*). The energy of the MKK6GRA-p38α and MKK6-p38α systems was minimized using the steepest descent integrator, and the solvated systems were equilibrated afterwards in the canonical (NVT) ensemble for 10 ns, with initial velocities sampled from the Boltzmann distribution at 310 K and a 2 fs timestep. The temperature was kept constant at 310 K by a velocity-rescale thermostat (*95*). The long-range electrostatics were calculated by the particle mesh Ewald algorithm, with Fourier spacing of 0.16 nm, combined with a switching function at 1.0 nm for the electrostatic and VdW interactions. The systems were then equilibrated for additional 10 ns in the isothermal-isobaric (NPT) ensemble prior to the production runs applying position constraints only to the protein (with a restraint spring constant of 1000 kJ mol^-1^ nm^-2^), where the pressure was maintained at 1 bar through a cell-rescale barostat (*96*).

Post equilibration, we ran a series of independent production runs in the NPT ensemble, coupled with a velocity-rescale thermostat (*89*) at 310 K and a cell-rescale barostat (*96*) at 1 bar for each of the systems as detailed in Table S6. Both unrestrained and restrained sets of replicas for MKK6GRA-p38α are randomly initialized from the same initial conformation and differ by the application of a harmonic restraint on the distance to the native inter-kinase contact found at the interface of the two monomers to maintain the initial relative position of p38α and MKK6GRA. The distance to the native contact map as implemented in the PLUMED plug-in was restrained using a force constant of 1,500 kJ mol^-1^ nm^-2^. Since the movement of the A-loop of p38α was important, contacts that involved residues of the A-loop were removed from the final contact map. The definition of the atoms involved in each contact, as well as the parameters for the contact map can be found in the input files deposited in the YARETA repository (doi: 10.26037/yareta:jmnbt6lzkrfrpdcf3m3343smxq). (*97*)(*97*)(*101*)(*100*) We expanded the dataset of unrestrained simulations for MKK6GRA-p38α adding a set of three replicas starting from a conformation that originates from a snapshot of one of the restrained simulations in which the A-loop of p38α extended towards MKK6 ATP binding site and p38α Y182 was close to MKK6 ATP.

### Adaptive MD simulations setup

To better understand how MKK6 engages with p38α, we ran a series of adaptive MD simulations (*48*) that allowed us to extend our sampling of prebound and bound states. This approach has been used in the past to study the association kinetics of small proteins (*49*). To generate a series of starting conformations for the adaptive simulations, we started by running a 60 ns long adiabatic bias simulation (*98*) for each system (MKK6GRA-p38α, MKK6-p38α) to pull the centres of mass of the two kinases 6 nm apart. In these simulations, a time-dependent harmonic biasing potential with a force constant of 5,000 kJ mol^-1^ nm^-2^ on the centre of mass distance was applied to pull the two kinases away from the conformation seen in the cryo-EM structure. During the course of these simulations, we also applied a harmonic constraint with a force constant of 1,500 kJ mol^-1^ nm^-2^ on a contact map of the KIM with p38α to prevent KIM from unbinding from p38α after the breaking of the N-to-N and C-to-C lobe interfaces. At the end of the adiabatic bias simulations, we manually selected eight conformations for each system representing intermediate states of the detachment process. These conformations were then fed into the adaptive sampling pipeline.

The adaptive method for the MD simulations we ran was based on the following scheme using a GROMACS implementation of the adaptive algorithm found in HTMD (*99*). We subjected each of the starting conformations to an independent 50 ns long unbiased simulation. Then, we retrieved the simulations and analysed them using a Markov state model. From this analysis, we extracted a new set of starting conformations, which were used for the next epoch of parallel simulations, etc. The method with which the conformations for each epoch are selected is the key to speeding up the sampling. After the first epoch, the next epochs use the Markov model to select the next starting conformations. This approach favours the exploration of new and undersampled states. This strategy can yield large speed-ups compared to non-adaptive MD runs because slow processes with large energy barriers are broken down into steps with individually smaller barriers that can be sampled efficiently. Although the individual simulations are not started from an equilibrium distribution and are shorter than binding and dissociation timescales, Markov modelling approaches can combine such a trajectory ensemble towards unbiased equilibrium models of the long-timescale kinetics and thermodynamics.

For the construction of the on-the-fly MSM during the adaptive simulations, different combinations of features were tested. Out of the tested feature combinations, the following features were used for the construction of the MSM that was used to span new simulations from: (i) the distance between the OG1 of p38α T180 and PG of MKK6 ATP, as a descriptor of the phosphorylation of T180, (ii) the distance between the OH of p38α Y182 and PG of MKK6 ATP, as a descriptor of the phosphorylation of Y182, (iii) the contact between p38α S32 and MKK6 N89, as a descriptor of the N-to-N lobe interaction, (iv) the contact between p38α Y258 and MKK6 K268, as a descriptor of the C-to-C lobe interaction, (v) the contact between p38α I116 and MKK6 L13, as a descriptor of the KIM-p38α interaction, and (vi) the contact between p38α F129 and MKK6 V7, as a descriptor of the KIM-p38α interaction. Contacts between residues were defined by nearest-neighbour heavy-atom distances <0.4 nm. For the construction of the MSM during the adaptive simulations, the three slowest linear combinations of the relevant contacts were computed variationally by the TICA method (*100*) (lagtime 20 ns), and this space was discretized using *k*-means clustering (*k*L=LJ500). Each epoch ran for 50 ns of unbiased simulation resulting in 9.2 μs to 9.6 μs of accumulated simulation time for the MKK6GRA-p38α, and MKK6-p38α, respectively (Table S8).

### Free energy simulations setup

To evaluate the relative stability of MKK6-p38α and MKK6 αG-helix mutant-p38α at the interaction region between MKK6 αG-helix and the hydrophobic pocket in p38α, we ran Multiple-Walkers (MW) metadynamics simulations in the well-tempered ensemble (*101*). Both systems were initialised from the cryo-EM conformation, and the distance with respect to the native contact map at the C-lobe interface and the distance between the centres of mass of p38α and MKK6 αG-helix (MKK6: residues 264-272, p38LJ: residues 228-240) were used as collective variables (CVs). The contact map was defined based on the resolved cryo-EM structure considering the sum of the contacts formed between each residue at the C lobes and its neighbouring ones within a radius of 0.5 nm. A force constant of 1,500 kJ mol^-1^ nm^-2^ was applied to the distance to the native contact map found at the interface of MKK6 KIM and p38α and at the N-to-N lobe interface to maintain the initial relative position of the two monomers while biasing the detachment at the C-to-C lobe interface. Harmonic restraints were also added to the two CVs to help convergence. The metadynamics simulations were terminated when an extensive exploration of the CV space was achieved. We used 8 walkers in the NPT ensemble and accumulated a simulation time of 2.9 μs and 2.5 μs for MKK6=p38α and MKK6 αG-helix mutant-p38α systems, respectively. The free energy profiles were reconstructed using the instantaneous metadynamics bias shifted by the c(t) factor, and block analysis was performed to compute the statistical error. The simulation parameters used in the metadynamics simulations were the same as the ones used in the unbiased MD simulations.

### MD simulations derived structures and analysis

To quantify the rotation of p38α with respect to MKK6 over the course of the simulations, we started by clustering the conformations derived from each MD simulation run. All the conformations visited during each simulation were clustered using the *gromos* algorithm (*102*) and the *g_cluster* routine (GROMACS) by using the RMSD of the backbone atoms of MKK6 as the distance between the structures and the cut-off value from 2 to 4.5 Å to ensure a representative number of frames in each cluster. The central structures of the most populated clusters of each basin (i.e. the structure with the smallest RMSD distance to all the other members of the cluster) were chosen as the representative ones of each simulation. The central structures of each simulation were then superimposed to the cryo-EM structure using the MKK6 as a reference and the rotation matrix of p38α was calculated through the ChimeraX functionalities.

For the kinetic-based clustering, we used all the simulations of each system where no constraints were applied (Table S8) since such constraints could slow down or accelerate certain transitions and affect the population of the resulting clusters. The MD data of each system was projected on the same feature space as the one used during the adaptive sampling (see above). The dominant four time-lagged independent (TICA) components, which represent the most slowly changing collective coordinates in the feature space were then calculated (lagtime 35 ns). The reduced space was used for cluster discretization into microstates with *k-*means clustering (*k*=500). Using the PCCA+ spectral clustering algorithm (*103*), we built a coarse-grained kinetic model by optimally grouping together the fast-mixing microstates resulting from the *k*-means clustering into metastable states.

Since the conformational ensembles that we obtained from all the unbiased MD simulations prior to any clustering are broad and the experimental data often have low information content and may be noisy, we used a Bayesian inference method and the maximum entropy principle framework to refine computationally-derived structural ensembles through experiments. In this framework, an ensemble generated using a prior model is minimally modified to match the experimentally observed data better. In particular, here, we combined ensemble averaged experimental SAXS data with the ensemble of conformations that we obtained after the kinetic clustering with the aim to obtain a structural ensemble of the system, which agrees with the experimental data. We used an iterative Bayesian/Maximum Entropy (iBME) protocol (*50, 104*), which takes as input simulations that were generated without taking the experimental data into account, and subsequently updates them using statistical reweighting. The purpose of the reweighting is to derive a new set of weights for each configuration in a previously generated ensemble so that the reweighted ensemble satisfies the following two criteria: (i) it matches the experimental data better than the original ensemble and (ii) it achieves this improved agreement by a minimal perturbation of the original ensemble.

To do so, we first calculated the SAXS profiles of each conformer of the ensemble using the Pepsi-SAXS algorithm (*105*), keeping the scale and background parameters fixed (*I*(0) = 1, *cst* = 0) and and setting the δ *=3*.34 e/nm^3^ (1% with respect to the bulk solvent) and *r*_0_ = 1.681 Å. The chosen values of the δρ and r_0_ parameters have been shown to give results that approximate well experimentally observed values for most folded proteins (*50*). After oprimising the θ parameter (θ=80), we optimised the weights, *I*(0) and *cst* using iBME against the experimental SAXS curves (Fig. S2) and obtained the ensemble with the best χ^2^.

To better characterise the conformations seen in p38α-MKK6GRA state A (Fig. 4, Movie S6), we clustered the ensemble using the *k-*means clustering. We used the elbow method to identify the optimal number of clusters (*k*=3). We performed clustering based on the backbone RMSD of the MKK6GRA and p38α cores with respect to the cryo-EM structure (PDB-8A8M), and a conformation extacted from one of the unrestrained MD simulation, where p38α N-lobe is rotated and T180 approaches MKK6 ATP (see main text).

## Supplementary Text

### Unrestrained simulations of MKK6-p38α indicate that Y182 can reach MKK6 ATP

Since the resolved cryo-EM structure has a chimeric MKK6 with a modified KIM on its N-terminal tail adopted from GRA24, we reasoned that the simulations of the MKK6-p38α with the WT KIM of MKK6 could give us more information on the role of this motif that the cryo-EM structure alone could not provide. In the WT, the “electrostatic hook” of KIM is recognised by p38α and forms ion pairs with residues of the β7/β8 turn of p38α. Indeed, during the simulations, K8 and R9 of the WT sequence anchor the KIM to p38α by forming backbone and side-chain hydrogen bonds and salt-bridges with E161, while the salt-bridge between the side chains of K14 and E160 further solidifies the KIM around p38α (Fig. S15). Unlike the set of simulations with the chimeric MKK6, in two out of the five simulations of this setup, we see both T180 and Y182 coming relatively close to MKK6 ATP (Table S9) without p38α having to rotate. Both during the approaching of the A-loop of p38α to the MKK6 ATP and after, the two kinases maintain their initial relative orientation throughout the simulations.

During the simulations of p38α with MKK6 comprising of the WT sequence, we were not able to see the rotation of p38α even though T180 got relatively close to ATP in one of the replicas. We think that the simulation time may not be sufficient for the N-lobe of p38α to rotate.

### Restraining the MKK6GRA-p38α relative position enhances the phosphorylation at Y182

We ran a set of five simulations for the MKK6GRA-p38α dimer, starting from the cryo-EM structure and applying mild harmonic restraints on the interface interactions to stabilise the relative orientation of the monomers in the complex (see Methods). Due to the presence of the restraints, we did not observe any large conformational change akin to the one seen in the unrestrained simulation described in the Main Text, showing T180 clearly interacting with MKK6 ATP.

In this set of restrained simulations, two out of the five replicas show p38α A-loop adopting an extended conformation with Y182 approaching P_γ_ of MKK6 ATP up to a final distance of 4.4 Å and 3.5 Å, respectively. In both cases, p38α ATP binding site slightly opens allowing the adenine group of p38α ATP to move out of the binding site. Interestingly, in one of the two simulations, we also see that the extended A-loop forms a one-turn helix (residues E178-Y182), which involves both phosphorylation sites. A similar transient helix on the middle section of the A-loop, which exposes the phosphorylation sites, has been reported in MD simulations of the unphosphorylated kinase domain of EGFR (*106, 107*), and is also reminiscent of a helix in the A-loop of MPSK1 kinase (*108*) (PDB-2BUJ). In the rest of the replicas of this set of simulations, the conformation adopted by the A-loop is not compatible with phosphorylation on either T180 or Y182 since the A-loop extends towards the p38α C-lobe or the MKK6 αC-helix.

It is worth mentioning that in all restrained simulations, the KIM never detaches from p38α regardless of the relative conformation – catalytically compatible or not – adopted by the two kinases.

### Released simulations of MKK6GRA-p38α confirm the role of MKK6 KIM in stabilising the active complex

The results obtained from the set of restrained simulations suggest a role of MKK6 KIM in stabilizing the MKK6-p38α complex. However, due to the presence of the harmonic restraints on the interface contacts, we were not able to evaluate to what extent MKK6 KIM contributes to the rotation of the p38α N-lobe.

We thus ran an additional set of three simulations releasing the restraints on the interface. We started the simulations from a representative conformation of the complex extracted from one of the restrained simulations showing Y182 pointing towards the MKK6 ATP (Y182-P_γ_=5.1 Å) and residues E178-Y182 of the A-loop folded into a one-turn helix.

In two out of three replicas the KIM remains stably in contact with p38α throughout the simulation.

Concurrently, we observe that in one simulation, p38α A-loop extends above the MKK6 αC-helix in a way that allows Y182 to interact with Q93, forming, thus, a catalytically incompatible complex. In the simulation we extended up to 2 μs, we observed the unfolding of the one-turn helix of the A-loop and T180 approaching MKK6 ATP. After clustering of this trajectory (RMSD cut-off distance equal to 2.75 Å), in the most populated cluster (cluster 1), Y182 and T180 fluctuate at 1-2 nm away from MKK6 ATP while in the second most populated cluster (cluster 2) they reach at a distance <1 nm. Interestingly, in cluster 1, the p38α N-lobe is turned by approximately 51° with respect to the initial conformation, while in cluster 2, in which both Y182 and T180 are closer to MKK6 ATP, the rotation is more significant, i.e., approximately 66°. This observation is consistent with what we observed with the unrestrained simulation showing the N-lobe having to rotate for T180 to interact with MKK6 ATP. Although interesting, from this simulation we were not able to confirm the role of the salt bridge between MKK6 K17 and p38α E160 in priming the rotation of p38α.

Lastly, in the third simulation of this set, we observed p38α A-loop moving from an extended to sequestered conformation concurrently with the dissociation of MKK6 KIM, thus confirming the role of MKK6 KIM in stabilizing the active complex. The conformational change of the A-loop required approximately 200 ns after the detachment of the KIM. Finally, even when most interface contacts are lost, the contacts around the hydrophobic patch between the C-lobes of p38α and MKK6 are maintained. Comparing the complex conformation with a reference structure for inactive p38α (PDB-3S3I) we were able to assess the inactive-like conformation of the A-loop adopted at the end of the simulation (RMSD ^p38a^ = 3.5 Å with respect to the crystal structure, Fig. S10). The KIM is also in contact with the hinge region of p38α. It is worth mentioning that during the simulations that we ran, we see that fluctuations of MKK6 are propagated through its N-terminal loop and KIM to the hinge region of p38α, affecting the stability of p38α ATP inside its pocket.

### p38LJ-MKK6 LJG-helix mutant populates conformations showing a detached C-to-C-lobe interface

We performed multiple-walkers metadynamics simulations to assess the relative stability of p38α in complex with MKK6 and MKK6 αG-helix mutant. Specifically, we biased the distance between the centres of mass of p38α and MKK6 αG-helix to enhance the separation of the complex and the distance to the native contact map at the C-to-C-lobe interface. By reconstructing the free energy surface underlying the detachment of the complex at the C-lobe interface, we were able to quantify the relative stability of the two complexes. In Fig. S9C, we reported the free energy profiles as a function of the distance to the native contact map of the cryo-EM structure at the C-lobe interface for both systems. The free energy surface of p38α-MKK6 αG-helix mutant indicates that the detached conformation at the C-to-C-lobe interface (contact map distance > 20 a.u.) is more populated than the conformation observed in the cryo-EM structure (Fig. S9D). Conversely, the p38α-MKK6 complex populates conformations showing the C-lobes interacting with each other at the hydrophobic patch as seen in the cryo-EM structure (contact map distance < 20 a.u., Fig. S9D).

These results agree with the biochemical data (Fig. 2F) and the HDX data (Fig. S9B), which show reduced activity in the presence of the αG-helix mutations and no shielding of p38α in the MAPK insert region by the MKK6^DD^ αG-helix mutant.

### Kinetic clustering of the p38α-MKK6 WT reveal intermediate states

Similar to the p38α-MKK6GRA, one of the clusters of p38α-MKK6 corresponds to the face-to-face conformation, and the A-loop of p38α adopts a conformation that brings T180 and Y182 close to the MKK6 ATP (state F, Movie S11). However, unlike the p38α-MKK6GRA, we do not see the rotated p38α in this ensemble. The second macrostate is again similar to state B of p38α-MKK6GRA in which the p38α A-loop has a conformation similar to the one seen in the cryo-EM structure, but the N-to-N lobe interaction is lost (state G, Movie S12). Finally, apart from the state that corresponds to the non-specific binding of MKK6 (state H, Movie S13), we see again a late intermediate state in which the KIM is bound to its recognition site while the N-lobes are separated (state J, Movie S14), reminiscent of a tightly bound state. Figure S12 displays all the macrostates that were identified for the p38α-MKK6.

### Conformations seen within the face-to-face macrostate of p38α-MKK6GRA

<u>Clustering</u> of the conformations in the catalytically relevant state A of p38α-MKK6GRA (Fig. 4, Move S6) showed the presence of three distinct conformations. In particular, these conformations correspond to the face-to-face cryo-EM structure, the N-lobe rotated p38α, and an intermediate conformation where the two kinases are held together by the C-to-C lobe interface and the N-lobe of p38α is not fully rotated yet.

## Acknowledgments

PJ was supported by a EMBL predoctoral fellowship. We thank Stephen Cusack and Andrew McCarthy (EMBL Grenoble) for encouragement, Wojtek Galej (EMBL Grenoble) for critical reading of the manuscript and Mohamed-Ali Hakimi (IAB, Grenoble) for introducing us to the *T. gondii* protein GRA24. We acknowledge Sarah Schneider for her support in using the EM Facility at EMBL Grenoble and thank Felix Weiss, Wim Hagen and Simon Fromm for excellent data collection at the EMBL Imaging Centre. We thank Alice Aubert (EMBL, Grenoble) for support in eukaryotic protein expression, the proteomic core facility at EMBL Heidelberg for mass spectrometry and Ulrike Kapp (ESRF) for assistance in the crystallisation of p38α. This work used the platforms of the Grenoble Instruct-ERIC centre (ISBG ; UAR 3518 CNRS-CEA-UGA-EMBL) within the Grenoble Partnership for Structural Biology (PSB), supported by FRISBI (ANR-10-INBS-0005-02) and GRAL, financed within the University Grenoble Alpes graduate school (Ecoles Universitaires de Recherche) CBH-EUR-GS (ANR-17-EURE-0003). We acknowledge the European Synchrotron Radiation Facility for the provision of synchrotron radiation facilities, and we would like to thank the staff of the ESRF and EMBL Grenoble for assistance and support in using beamlines ID23-1 and ID23-2. SAXS data were collected at beamline B21 at Diamond Light Source, UK. We acknowledge Massimiliano Bonomi (Institut Pasteur - CNRS, France) for discussions and technical support in cross-correlation analysis. Molecular graphics and analyses were performed with UCSF ChimeraX, developed by the Resource for Biocomputing, Visualization, and Informatics at the University of California, San Francisco, with support from National Institutes of Health R01-GM129325 and the Office of Cyber Infrastructure and Computational Biology, National Institute of Allergy and Infectious Diseases. We acknowledge PRACE and the Swiss National Supercomputing Centre (CSCS) for large supercomputer time allocations on Piz Daint, project IDs: pr126, s1107.

## Funding

EMBL (MWB)

Swiss National Science Foundation [projects number: 200021_204795] (FLG).

Bridge [40B2-0_203628] (FLG).

## Author contributions

Conceptualization: PJ, EP, FLG and MWB

Methodology: PJ, IG, DG, JvV, MP and OV

Investigation: PJ, IG, DG, JvV, OV, MT, FLG, EP, and MWB

Visualization: PJ, IG, DG, JvV and OV

Funding acquisition: FLG and MWB

Supervision: FLG, EP and MWB

Writing - original draft– PJ and MWB

Writing – review & editing: PJ, IG, DG, JvV, MP, OV, MT, FLG, EP, and MWB

## Data and materials availability

Accession codes: Structure of the MKK6^DD^GRA-p38α^T180V^complex: EMPIAR-11203 (raw micrographs; Electron Microscopy Public Image Archive), EMD-15233 (map; Electron Microscopy Data Bank) and PDB-8A8M (coordinates; Protein Data Bank). Structures of apo p38α and p38α^K53R^: PDB-5ETC and PDB-5ETI (coordinates and structure factors; Protein Data Bank). The SAXS curves and pair distribution functions are deposited in the Small Angle Scattering Biological Data Bank (access codes: SASBDB-SASDRC6 and SASBDB-SASDRD6). The mass spectrometry proteomics data have been deposited to the ProteomeXchange Consortium via the PRIDE partner repository with the dataset identifier PXD040499 and 10.6019/PXD040499. The MD simulations data (trajectories, topology files, and structures of the SAXS-refined ensembles) are deposited in the YARETA repository (doi: 10.26037/yareta:jmnbt6lzkrfrpdcf3m3343smxq).

---

**Fig. S1.**
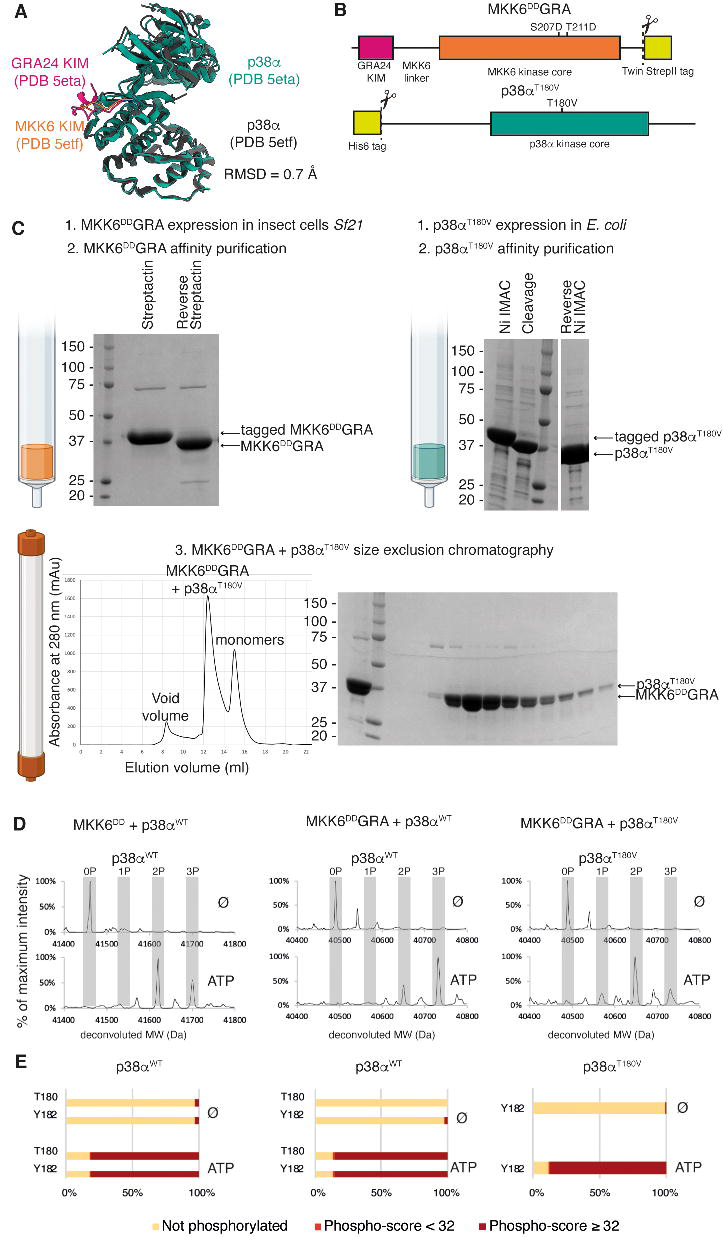
Expression, purification and activity of the MKK6^DD^GRA-p38α^T180V^ complex. (**A**) The *Toxoplasma* GRA24 KIM induces the same conformational change on p38α as the native MKK6 KIM, despite having higher affinity. The RMSD between the MKK6 KIM – p38α crystal structure (PDB 5etf) and the GRA24 KIM – p38α crystal structure (PDB 5eta) is 0.7 Å. **(B)** Constructs of MKK6^DD^GRA and p38α^T180V^used for structural studies. (**C**) MKK6^DD^GRA and p38α^T180V^were respectively expressed in *Sf21* and *E. coli*. They were purified by affinity purification (StrepTactin and Ni IMAC, respectively), followed by cleavage with 3C protease, and reverse affinity purification. Both proteins were mixed, and the complex was purified with size-exclusion chromatography. (**D-E**) Mass spectrometry (Native ESI-QTOF in (**D**) and LC-MS/MS in (**E**) reveals that the MKK6^DD^GRA chimera can phosphorylate p38α^WT^ as the MKK6^DD^ construct, as well as p38α^T180V^. In the presence of ATP, several sites on both proteins are phosphorylated, including both T180 and Y182 residues of p38α^WT^ A-loop and Y182 residue of p38α^T180V^ A-loop.

**Fig. S2.**
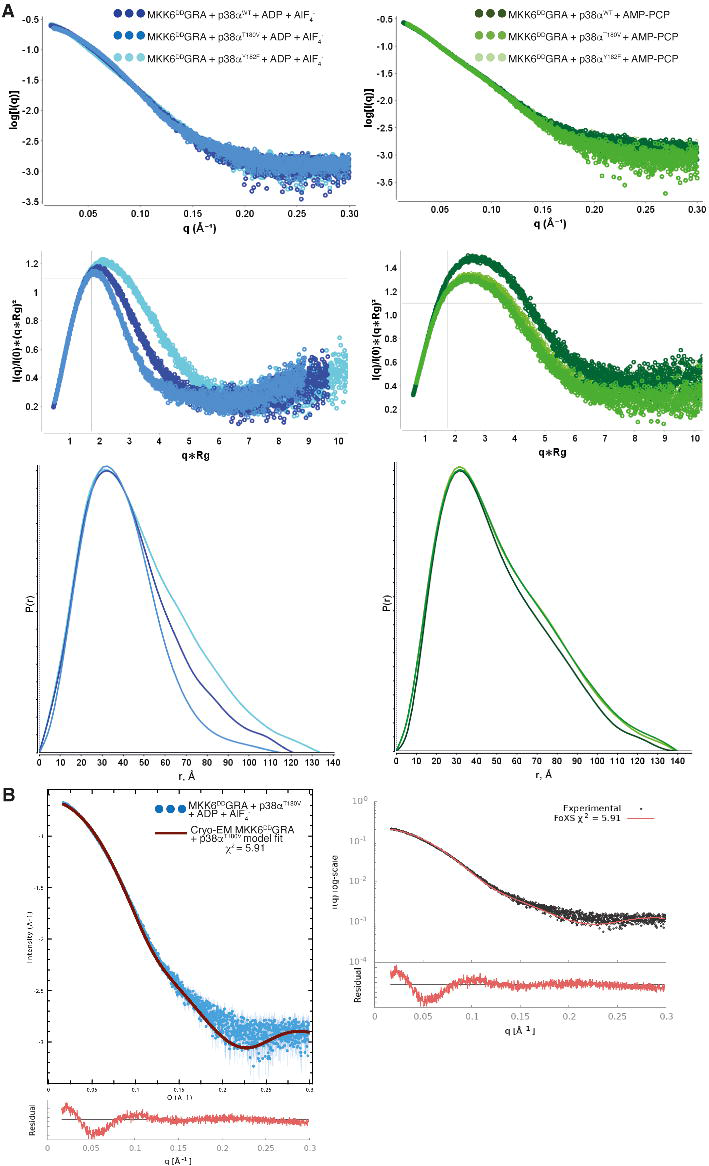
SAXS curves for MKK6^DD^GRA + p38α mutant complexes. (**A**) The blue panels show the IvsQ, Kratky, and P(r) plots for MKK6^DD^GRA + p38α mutants + ADP + AlF_4_^−^ complexes. The complex with p38α^T180V^ mutant is more compact than the one with p38α^WT^, which is more compact than the one with p38α^Y182F^mutation. The green panels are the same plots for the MKK6^DD^GRA + p38α mutants + AMP-PCP complexes. The complex with p38α^WT^ has a slightly increased Rg compared to the two mutant ones, but all share a similar D_max_. (**B**) χ^2^ fits for the cryo-EM MKK6^DD^GRA + p38α^T180V^ model (brick).

**Fig. S3.**
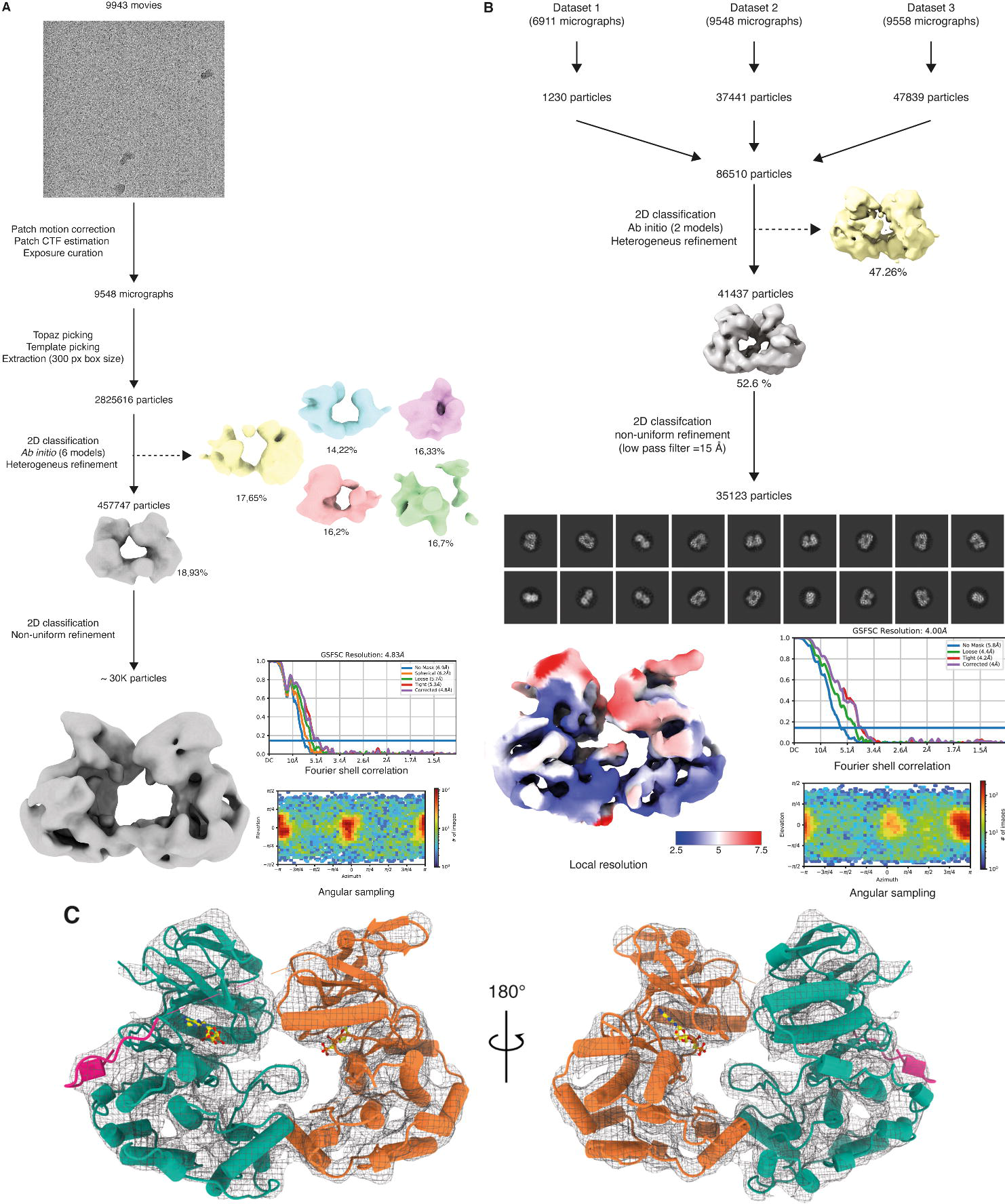
Cryo-EM processing and data assessment of the MKK6^DD^GRA-p38α^T180V^ model. (**A**) Exemplary workflow processing for one of the three datasets collected (dataset 2). Data processing was performed in cryoSPARC. A representative micrograph, typical result from heterogeneous refinement, final map with angular sampling and gold-standard Fourier Shell Correlation are also shown. (**B**) Cryo-EM processing of the final particle selection from the three merged datasets to obtain the final reconstruction at 4.0 Å resolution. The final sharpened map, coloured according to local resolution, angular sampling and gold-standard Fourier Shell Correlation are also shown. Representative 2D classes, which yielded the final map, are also shown. (**C**) Fitting of the model into the cryo-EM map. The MKK6^DD^GRA-p38α^T180V^ complex is shown in cartoon representation with α-helices as cylinders, and the sharpened map is shown as a black mesh.

**Fig. S4.**
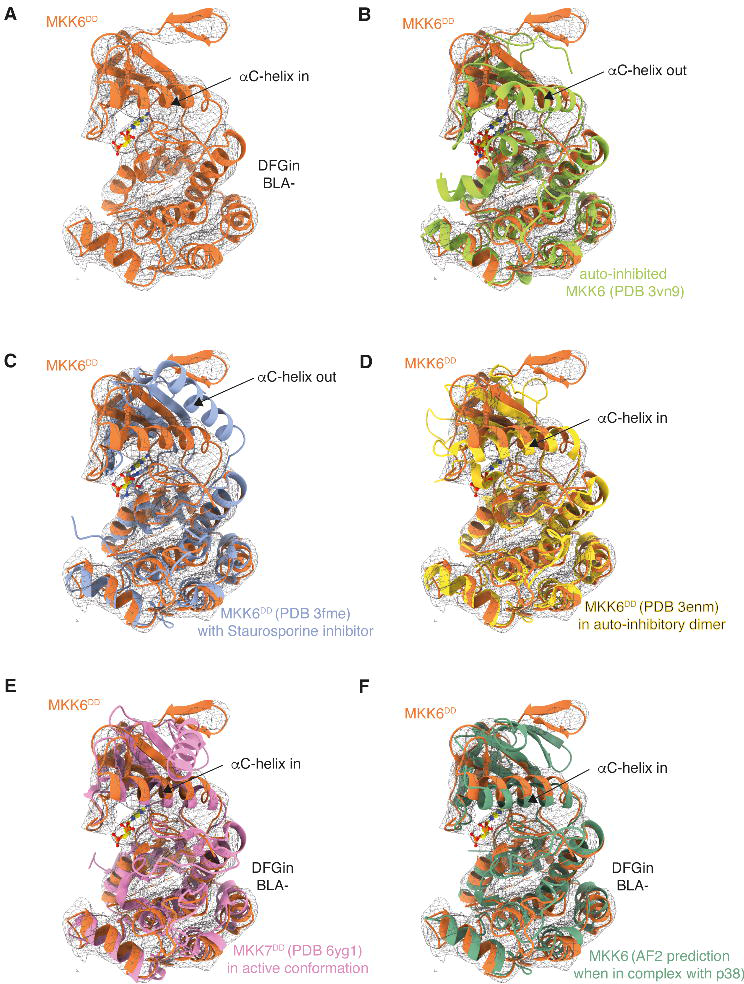
Comparison of MKK6 models with the cryo-EM structure. (**A**) MKK6^DD^ cryo-EM model fitted into the sharpened cryo-EM map (shown as a black mesh). (**B-F**) Overlay of the cryo-EM structure of MKK6^DD^GRA with (**B**) the crystal structure of MKK6 in an inactive conformation with AMP-PNP bound, (**C**) the crystal structure of MKK6 in an inactive conformation inhibited with staurosporin, (**D**) the crystal structure of MKK6 in an auto-inhibitory dimer, (**E**) the crystal structure of active MKK7 and (**F**) the AlphaFold2 model of MKK6 when in complex with p38α.

**Fig. S5.**
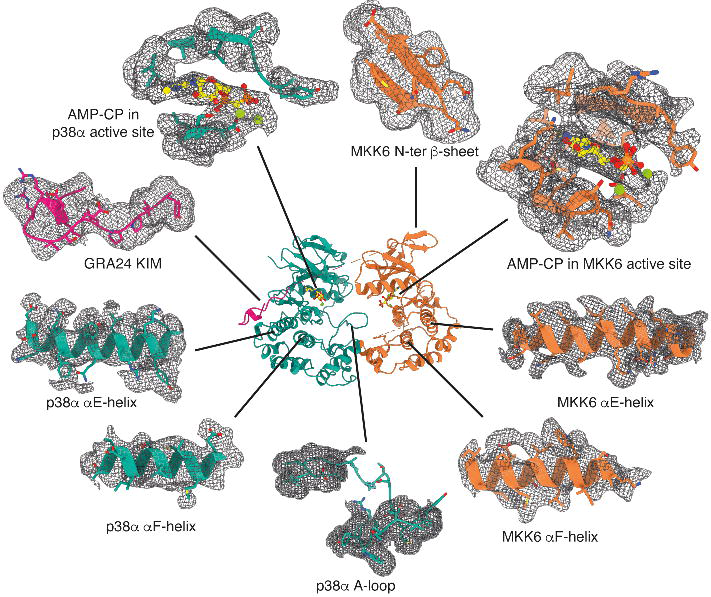
Detailed views of the MKK6^DD^GRA-p38α^T180V^ model with the sharpened Coulomb potential map. Structural elements of the model are shown in cartoon and stick representation and the sharpened map is shown as a black mesh.

**Fig. S6.**
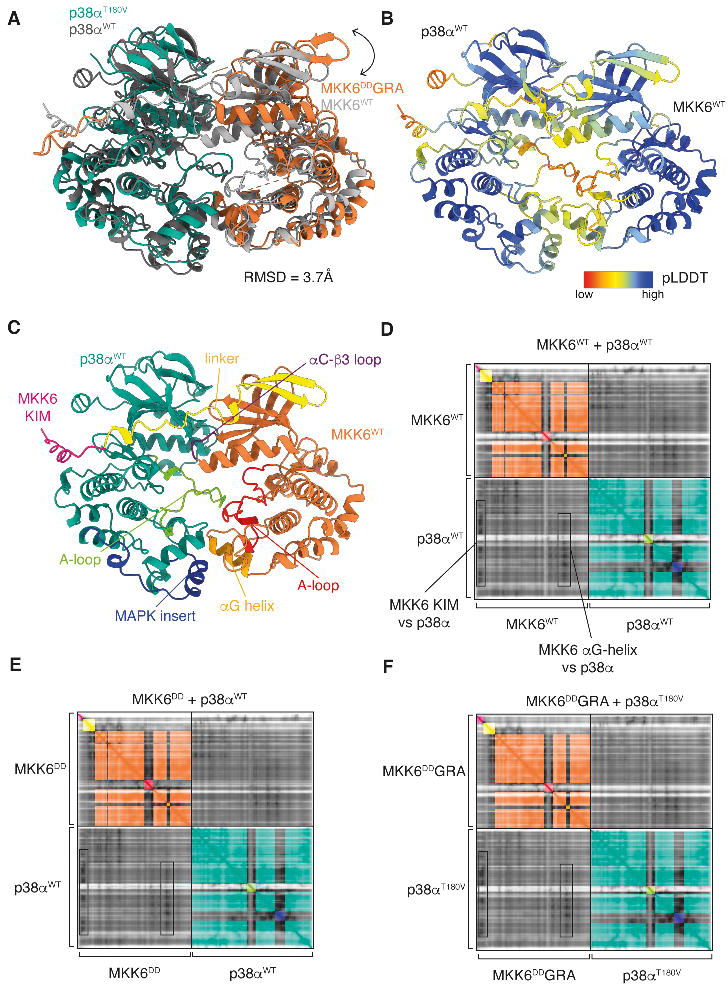
AlphaFold2 multimer predicts a similar interface of the complex. (**A**) MKK6^DD^GRA-p38α^T180V^ model (colour), superimposed with the AlphaFold2 prediction of the complex (grey). Differences in domain orientation are indicated by arrows and the overall RMSD value of the superposition is indicated. (**B**) Cartoon representation of the MKK6-p38α predicted model, coloured according to its per-residue confidence (pLDDT). (**C**) AlphaFold2 predicted MKK6-p38α model with relevant motifs indicated. (**D**) Predicted Aligned Error (PAE) plots for the MKK6-p38α prediction. Low PAE values (0 to 30 Å, decrease in grey scale) indicate higher confidence in respective residue placement. Within protein domains, PAE values are lowest, however, intermolecular placement, seen in the bottom left and top right quadrants, show confident placement, indicated by dark grey. For orientation, domains and relevant motifs in the plot are coloured as in (**C**). (**E**-**F**) PAE plots for MKK6^DD^GRA-p38α and MKK6^DD^GRA-p38α^T180V^ predictions, respectively, showing similar PAE values at the interfaces between the kinases.

**Fig. S7.**
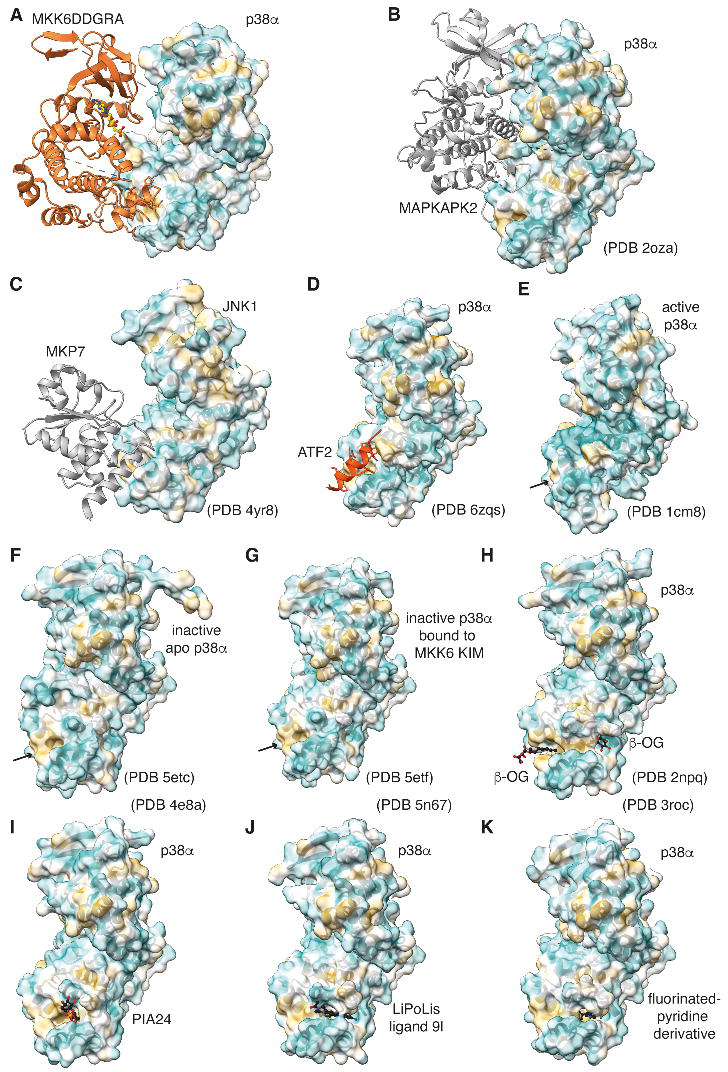
Interaction with, and variability of, the p38α hydrophobic pocket. The p38α surface is coloured according to hydrophobicity. (**A**) Cryo-EM MKK6^DD^GRA-p38α^T180V^model. (**B**) Crystal structure of p38α bound to its substrate MAPKAPK2 in an inactive complex showing a different binding mode. (**C)** Crystal structure of JNK1 bound to MKP7 phosphatase showing a similar binding mode as the MKK6 αG helix. (**D)** Crystal structure of p38α bound to substrate ATF2 peptide showing a similar binding mode as the MKK6 αG helix. (**E-G**) The p38α hydrophobic pocket is flexible: it is opened in inactive p38α crystal structures and closes in active p38α conformation. (**H-K**) The p38α hydrophobic pocket can bind small molecules and regulatory lipids: (**H**) β-octyl-D-glucopyranoside (β-OG) detergent, (**I**) phosphatidylinositol ether lipid analogue 24 (PIA24), (**J**) LiPoLis ligand 9l, and (**K**) a fluorinated-pyridine derivative.

**Fig. S8.**
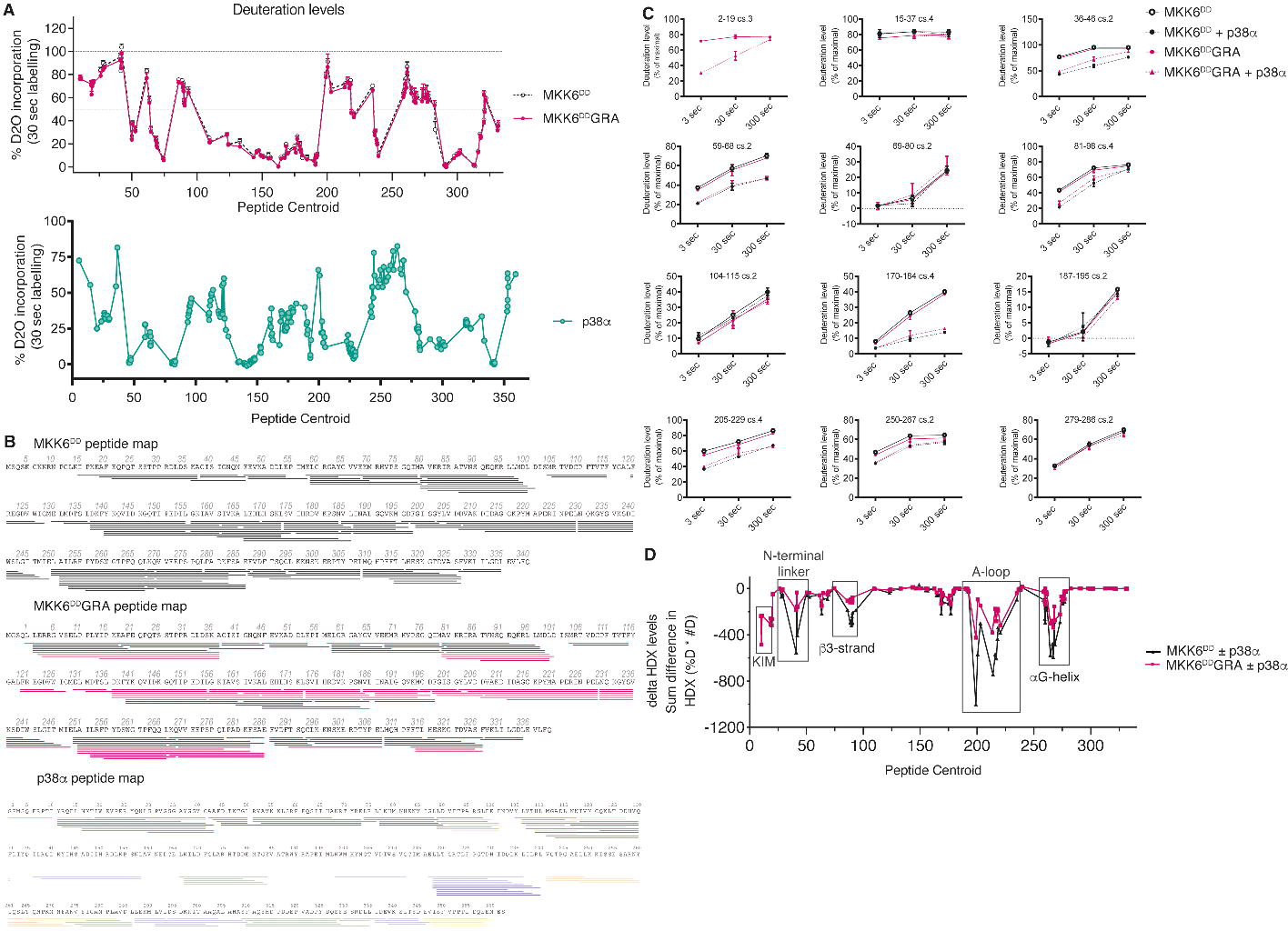
HDX-MS shows that MKK6^DD^GRA and MKK6^DD^ bind similarly to p38α. (**A**) Deuteration levels for each peptide of both MKK6^DD^ and MKK6^DD^GRA. Each dot represents one peptide plotted on the x-axis according to the central residue of the peptide (centroid). (**B**) Peptide map for both MKK6 proteins and p38α. Each line represents a high-quality peptide analysed in this study. (**C**) Uptake plots of a representative selection of peptides. Residue numbering based on MKK6^DD^. Data are mean +/− SD, n=3 (**D**) Graph plotting the difference in H/D exchange level for each studied peptide. Values indicate the sum of differences measured at three deuteration times (3, 30 and 300 sec), as the product of differences in % deuteration (%D) * the number of deuterons (#D).

**Fig. S9.**
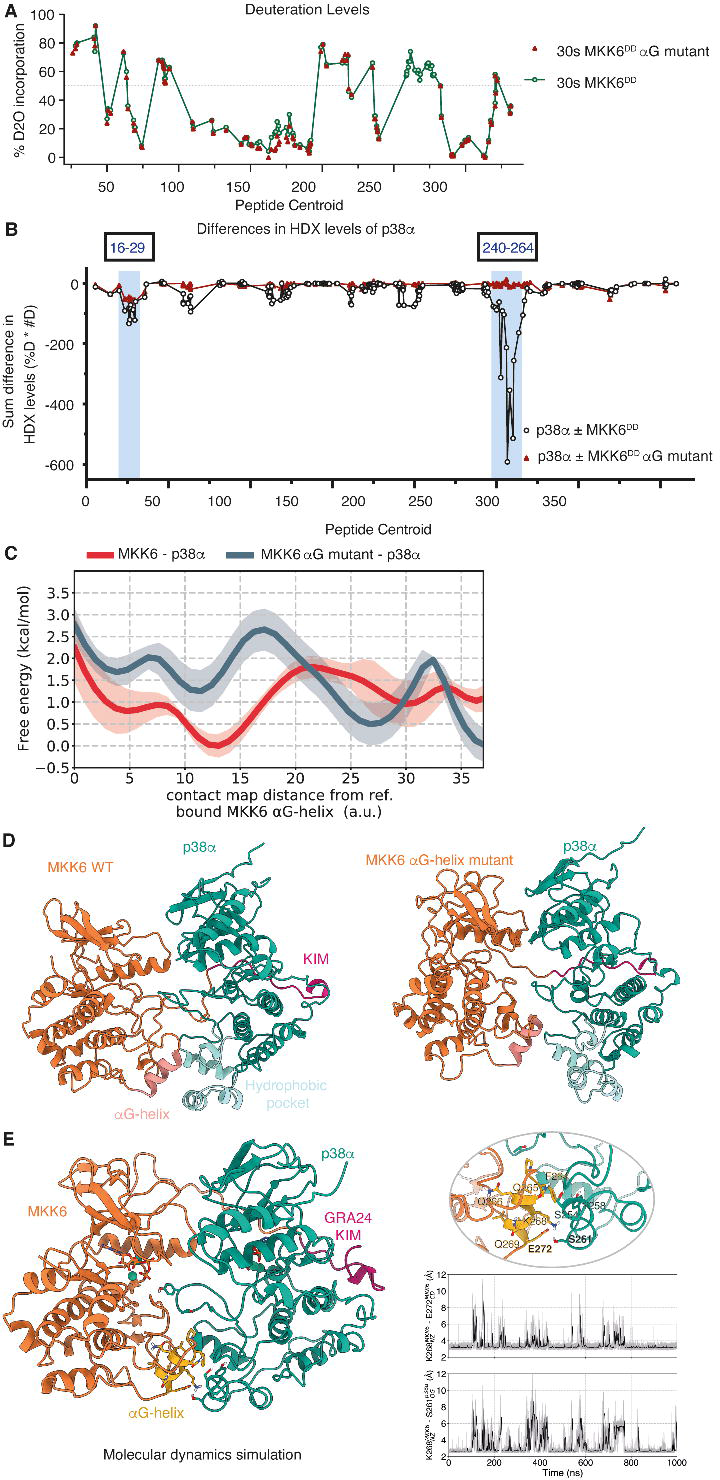
The MKK6αG-helix – p38α hydrophobic pocket interaction is stable and the MKK6^DD^ αG-helix mutant abolishes the interaction. (**A**) Deuteration levels for each peptide of both MKK6^DD^ and MKK6^DD^ αG-helix mutant. Each dot represents one peptide plotted on the x-axis according to the central residue of the peptide (centroid). (**B**) Two regions of p38α are protected from exchange upon MKK6^DD^ binding: residues 16-29 and residues 240-264, with the 240-264 region showing higher protection levels compared to the N-terminal region. Only the 16-29 region of p38α is protected from exchange upon MKK6^DD^αG-helix mutant binding. (**C**) p38α-MKK6 (red) and p38α-MKK6 αG-helix mutant (blue) free energy profiles as a function of the distance to the native contact map of the cryo-EM structure at the C-lobe interface. Shaded error bands show the estimates of uncertainty in the free energy in each region of the CV space. (D) Representative structures of p38α-MKK6 (left) and p38α-MKK6 αG-helix mutant (right) found in the free energy global minima are shown in cartoon representation. The C-to-C-lobe interface is maintained in p38α-MKK6. Conversely, the p38α-MKK6 αG-helix mutant populates conformations showing the detachment of MKK6^DD^ αG-helix from the p38α hydrophobic pocket. (**E**) Important interactions around the hydrophobic patch maintained over the course of the MD simulations, inset shows details of observed interactions and plots of distances during the simulation.

**Fig. S10.**
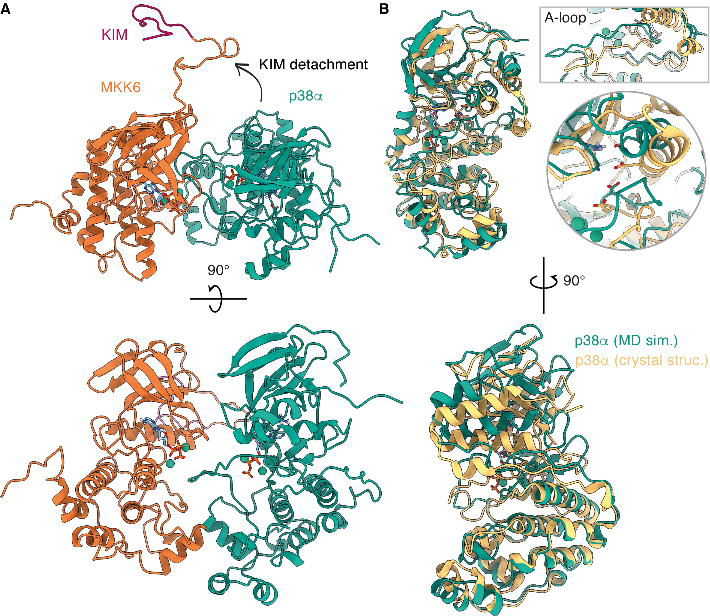
Communication between the MKK6 KIM and p38α. (**A**) Top and side views of the complex after the detachment of the KIM and consequent transition of p38α to an inactive-like conformation. (**B**) Superposition of p38α with the inactive p38α crystal structure (PDB 3s3i), RMSD^p38a^ = 3.5 Å. Detail: Top: A-loop that breaks the β-sheet motif normally seen in the active conformation and adopts an inactive-like conformation, Bottom: αC-helix and DFG motif and K/R53-E71 salt-bridge.

**Fig. S11.**
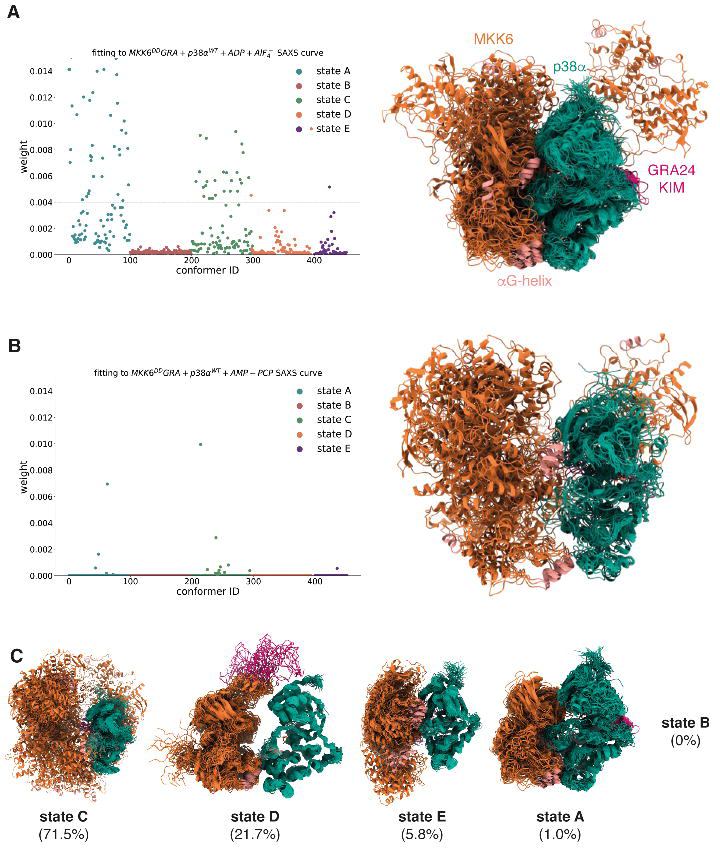
SAXS-based refinement of the MD-derived conformational ensemble through Bayesian/maximum entropy. (**A**) Statistical weights of the conformers after fitting (χ^2^=0.89) to the MKK6^DD^GRA + p38α^WT^ + ADP + AlF_4_ ^−^ SAXS curve (Fig. S2). Graphical representation of the conformations with a weight higher than 0.004. (**B**) Statistical weights of the conformers after fitting (χ^2^=2.1) to the MKK6^DD^GRA + p38α^WT^ + AMP-PCP SAXS curve (Fig. S2). Graphical representation of the conformations with a weight higher than 0.004. (**C**) Population of states as derived from the fitting to the MKK6^DD^GRA + p38α^WT^ + AMP-PCP SAXS curve.

**Fig. S12.**
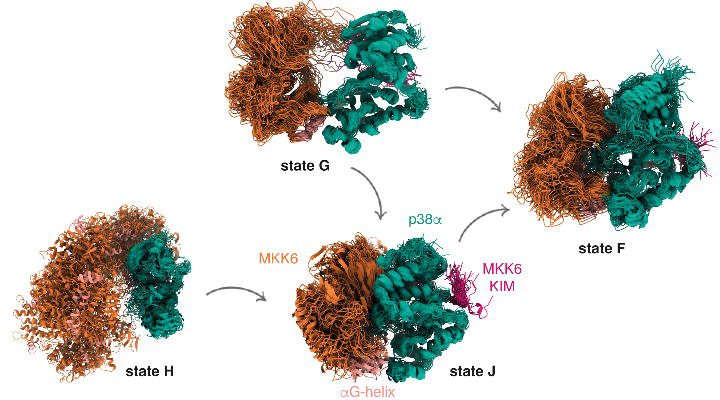
Kinetic-based clustering of the accumulated simulations of p38α-MKK6WT. Each macro-state shows an overlay of 50 representative conformations sampled proportionally to the equilibrium probability of each microstate in the corresponding macro-state and correspond to different kinetically distinct states.

**Fig. S13.**
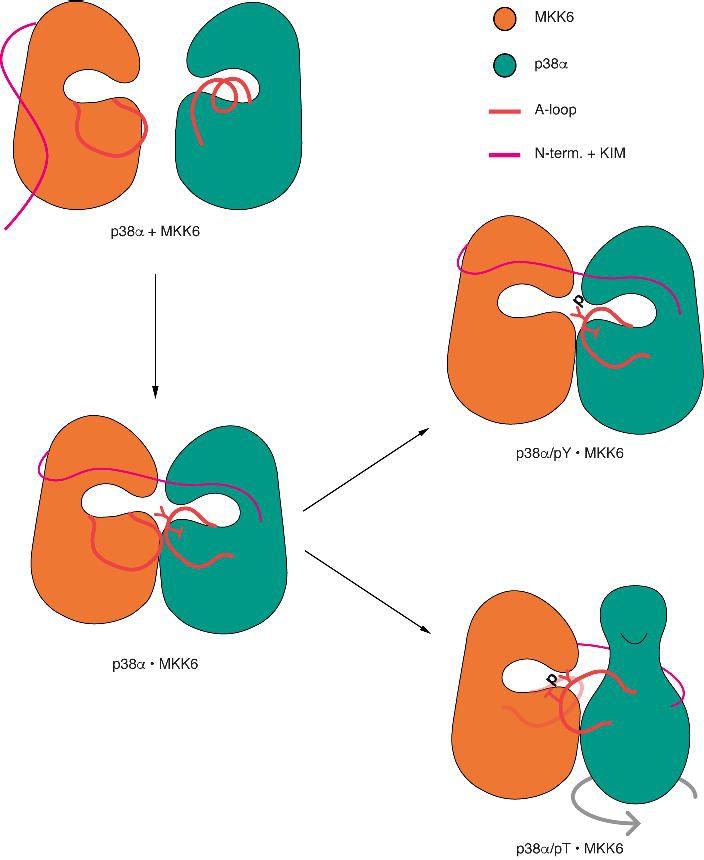
Schematic representation of the phosphorylation mechanism derived from the MD simulations. Although both p38α T180 and Y182 can reach MKK6 ATP at a catalytically competent distance, the N-lobe of p38α seems to have to rotate for T180 to reach MKK6 ATP in most replicas. The difference in experimentally derived kinetic rates measured for the first phosphorylation step may reflect the rotation of the N-lobe of p38α around its axis, which seems to be associated only with the T180 phosphorylation.

**Fig. S14.**
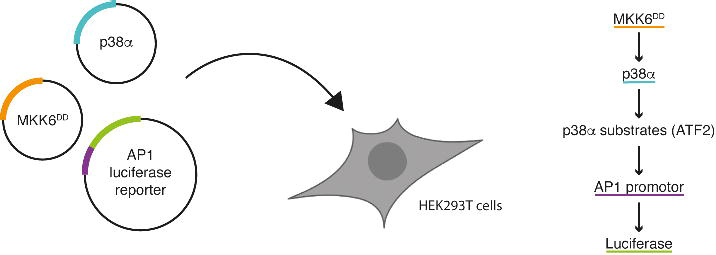
Cellular reporter assay. HEK293T cells are transiently transfected with MKK6^DD^ mutants (orange) and p38α (teal), together with a plasmid encoding firefly luciferase (green) under the control of the AP-1 promoter (purple). When p38α is activated, it phosphorylates several substrates, including ATF2, that will in turn stimulate the AP-1 promoter and trigger the expression of luciferase. Activity of p38α can be directly monitored by measurement of luminescence.

**Fig. S15.**
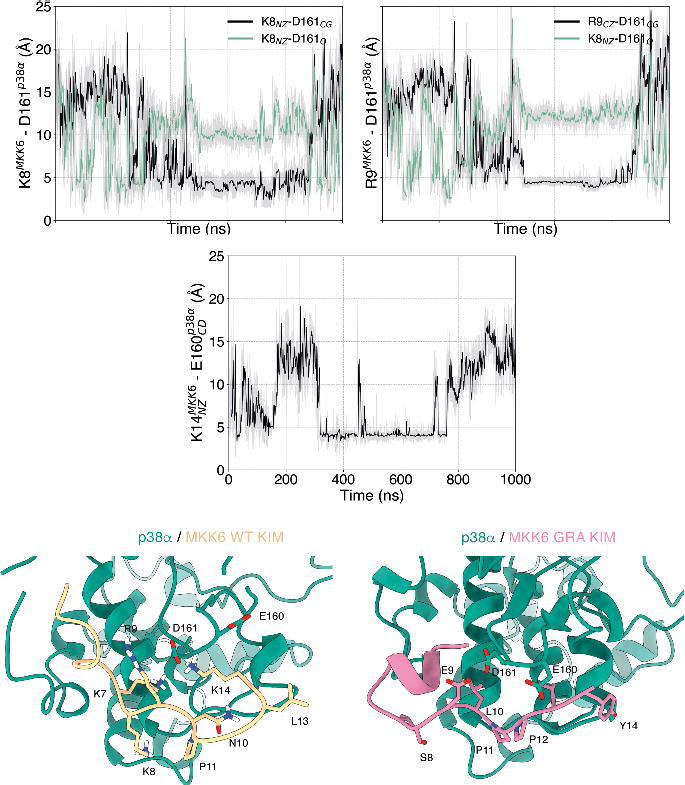
Important interactions that maintain the KIM of MKK6 in contact with p38α over the course of the MD simulations. The rolling average of the timeseries of side-chain-to-side-chain interactions is shown in black, while the side-chain-to-backbone interactions are shown in green. The reported timeseries refer to one of the simulations with the MKK6 WT sequence (Table S6), however, the same pattern of interactions is observed in three out of the five independent replicas where p38α stays in contact with MKK6.

**Table S1.** Calculated SAXS data using Scatter and FoxS.

| | I(0) | Rg (Å) | MW<br>estimation<br>(kDa) | D<sub>max</sub><br>(Å) | $\chi^2$ against<br>cryo-EM<br>MKK6<sup>DD</sup>GRA +<br>p38α<sup>T180V</sup> model | $\chi^2$ against MD<br>MKK6 + rotated<br>p38α<sup>WT</sup> model |
| --- | --- | --- | --- | --- | --- | --- |
| MKK6<sup>DD</sup>GRA + p38α<sup>WT</sup> +<br>AMP-PCP | 0.30 | 41.6 | 68 | 139 | 136 | 94.0 |
| MKK6<sup>DD</sup>GRA + p38α<sup>T180V</sup> +<br>AMP-PCP | 0.23 | 39.1 | 77 | 140 | 138 | 96.6 |
| MKK6<sup>DD</sup>GRA + p38α<sup>Y182F</sup> +<br>AMP-PCP | 0.30 | 39.4 | 70 | 139 | 165 | 115 |
| MKK6<sup>DD</sup>GRA + p38α<sup>WT</sup> +<br>ADP + AlF<sub>4</sub><sup>-</sup> | 0.26 | 32.2 | 65 | 121 | 19.1 | 10.1 |
| MKK6<sup>DD</sup>GRA + p38α<sup>T180V</sup> +<br>ADP + AlF<sub>4</sub><sup>-</sup> | 0.23 | 29.6 | 59 | 115 | 5.91 | 6.46 |
| MKK6<sup>DD</sup>GRA + p38α<sup>Y182F</sup> +<br>ADP + AlF<sub>4</sub><sup>-</sup> | 0.28 | 35.3 | 69 | 134 | 86.6 | 55.1 |
| cryo-EM MKK6<sup>DD</sup>GRA +<br>p38α<sup>T180V</sup> model | | 29* | | | | |
| MD MKK6 + rotated<br>p38α<sup>WT</sup> model | | 32* | | | | |

\* Theoretical Rg calculated using ATSAS

**Table S2.** X-ray data processing and refinement statistics.

| Structure | p38α (PDB-5ETC) | p38$\alpha^{K53R}$ (PDB-5ETI) |
| --- | --- | --- |
| Space group | $P2_12_12_1$ | $P2_12_12_1$ |
| Wavelength | 0.873 Å | 1.063 Å |
| Unit cell dimensions (Å) a, b, c | 45.1, 86.9, 124.0 | 45.4, 87.0, 122.9 |
| $\alpha, \beta, \gamma$ (°) | 90.0, 90.0, 90.0 | 90.0, 90.0, 90.0 |
| Resolution range (Å) | 20-2.42 | 20-2.8 |
| Number of unique reflections | 16774 | 10748 |
| Multiplicity<sup>1</sup> | 3.7 (2.8) | 3.9 (3.9) |
| Completeness<sup>1</sup> (%) | 87.3 (52.3) | 87.9 (90.8) |
| Rmerge<sup>1</sup> | 0.09 (0.50) | 0.17 (0.58) |
| $\langle I/\sigma(I) \rangle$<sup>1</sup> | 11.0 (1.9) | 8.3 (3.1) |
| Wilson B factor | 26 Å<sup>2</sup> | 28 Å<sup>2</sup> |
| No. of atoms | 3062 | 2918 |
| Protein | 2865 | 2867 |
| Ligands/ion | 5 | 0 |
| Water | 192 | 64 |
| R-factor<sup>3</sup> (%) | 19.4 | 17.8 |
| Free R-factor<sup>4</sup> (%) | 24.2 | 22.5 |
| RMS deviations: |  |  |
| Bonds (Å) | 0.013 | 0.008 |
| Angles (°) | 1.37 | 1.23 |

<sup>1</sup> Statistics for the highest resolution bin (2.95-2.8; 2.52-2.4) are shown in parenthesis

**Table S3.** Summary of cross-correlation scores for the rebuilt model and the predicted structures for different regions of each protein. The highest cross-correlation score is highlighted. The region with the highest cross-correlation score from each model was used in the final model.

| | Entire complex | MKK6-p38α hydrophobic patch<sup>1</sup> | MKK6 KIM domain<sup>2</sup> | p38α A-loop<sup>3</sup> | MKK6 region close to the linker<sup>4</sup> |
| --- | --- | --- | --- | --- | --- |
| Rebuilt model | 0.492 | 0.441 | 0.488 | 0.305 | 0.248 |
| AF2 model1 | 0.271 | 0.152 | 0.349 | 0.288 | 0.272 |
| AF2 model2 | 0.204 | 0.190 | 0.401 | 0.232 | 0.010 |
| MODELLER model1 | 0.477 | 0.497 | 0.450 | 0.307 | 0.239 |
| MODELLER model2 | 0.474 | 0.486 | 0.427 | 0.317 | 0.226 |
| MODELLER model3 | 0.462 | 0.478 | 0.436 | 0.320 | 0.140 |
| MODELLER model4 | 0.486 | 0.498 | 0.439 | 0.322 | 0.263 |
| MODELLER model5 | 0.479 | 0.396 | - | 0.315 | - |
| Conformation derived from MD simulations clustering | 0.283 | 0.242 | 0.351 | 0.261 | 0.174 |

<sup>1</sup> MKK6: res. 262-275, p38α : res. 252-262
<sup>2</sup> MKK6: res. 0-15, p38α : res. 108-136, 156-167, 309-318
<sup>3</sup> p38α : res. 172-189
<sup>4</sup> MKK6: res. 36-48

**Table S4.** Cryo-EM data collection, processing and refinement.

| **Data collection and processing** | **MKK6<sup>DD</sup>GRA-p38α<sup>T180V</sup> complex (PDB-8A8M)** |
| --- | --- |
| Magnification | 215,000 |
| Voltage (kV) | 300 |
| Electron exposure (e<sup>-</sup>/Å<sup>2</sup>) | 62.77 |
| Defocus range (μm) | -1.5 to -3 |
| Pixel size (Å) | 0.638 |
| Final particle images (no.) | 35,123 |
| Map resolution (Å) | 4.0 Å |
| FSC threshold | 0.143 |
| Map resolution range (Å) | 2.7-6.5 (25<sup>th</sup> to 75<sup>th</sup> percentile) |
| **Refinement** |  |
| Initial models used | 5ETA, 5ETC and a homology model of 6YG1 |
| Map sharpening factor (Å<sup>-2</sup>) | 200 |
| Model composition |  |
| Non-hydrogen atoms | 5249 |
| Protein residues | 5157 |
| Ligands | 92 |
| R.M.S deviations |  |
| Bond length (Å) | 0.004 |
| Bond angles (°) | 0.934 |
| Validation |  |
| MolProbity score | 2.44 |
| Clash score | 28.69 |
| Poor rotamers (%) | 0.52 |
| Ramachandran plot |  |
| Favoured (%) | 91.69 |
| Allowed (%) | 8.31 |
| Outliers (%) | 0 |

**Table S5. HDX results**

Provided in .xlsx file format

**Table S6.** Statistical analysis of cellular assay (Fig. 2)

| **p38α** | **MKK6** | **Mean** | **se** | **Shapiro**<br>**Wilk** | **Welch t test**<br>**against**<br>**p38α+MKK6<sup>DD</sup>** | **Significance**<br>**(<0.5 \*; <0.01**<br>**\*\*;**<br>**< 0.001 \*\*\*)** |
| --- | --- | --- | --- | --- | --- | --- |
| + | - | 2% | 0.002 | 4,36E-01 | 3,66E-12 | *** |
| + | DD<sup>1</sup> | 100 % | 0.057 | 7,71E-01 | 1,00E+00 | NS |
| + | αG-helix mutant<sup>2</sup> | 31 % | 0.023 | 7,76E-01 | 1,68E-10 | *** |
| + | β3-αC-loop<br>mutant<sup>3</sup> | 168 % | 0.122 | 8,21E-01 | 3,57E-04 | *** |

<sup>1</sup> S207D, T211D
<sup>2</sup> F264A, Q265A, L267A, K268A, E272A
<sup>3</sup> T87A, V88A, N89A

**Table S7.**
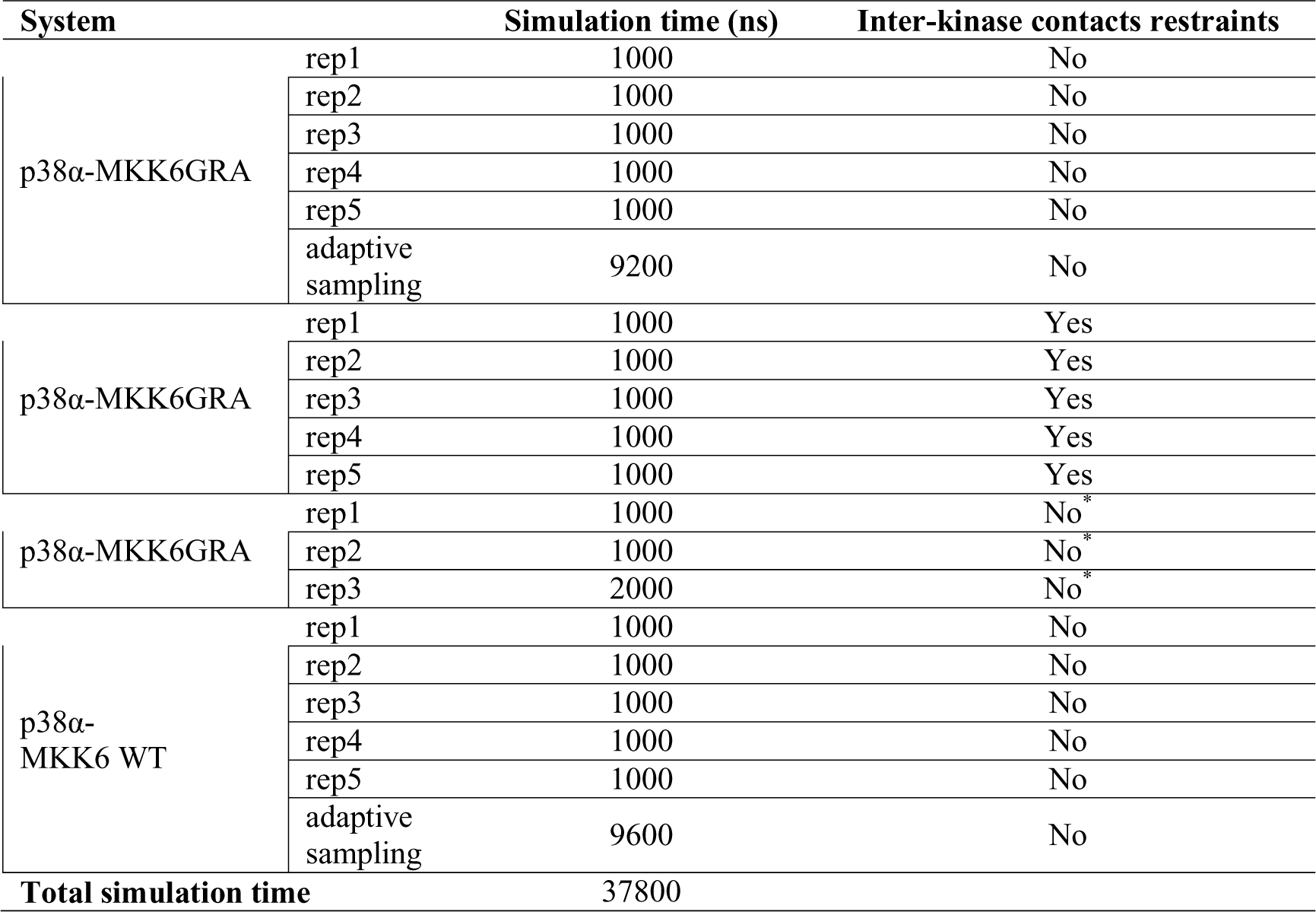
Summary of the unbiased simulations of the p38α-MKK6. The reported times correspond to the simulation time of each independent unbiased simulation. In the case of the adaptive sampling, we report the accumulated simulation time of all epochs. * started from a conformation originating from the restrained simulations of p38α-MKK6GRA, where p38α Y182 was close to MKK6 ATP.

**Table S8.** Comparison of MD-derived atom distances (Å) important for the phosphorylation of p38α at T180 and Y182 by MKK6 with atom distances reported for the SP20 phosphoryl transfer mechanism catalysed by the cAMP-dependent protein kinase (PKA) The reported distances for the PKA-SP20 phosphorylation were derived from QM/MM calculations (*46*), while the MD distances where derived from a frame where either T180 or Y182 was the closest to MKK6 ATP.

|  | QM/MM | MD simulations |  |  |  |
| --- | --- | --- | --- | --- | --- |
| | PKA-SP20 | MKK6GRA-p38α<br>(rep3 / rep5,<br>no restraints) | MKK6GRA-p38α<br>(rep3 / rep4,<br>with restraints) | MKK6GRA-p38α<br>(rep3,<br>released restraints) | MKK6-p38α<br>(rep2 / rep5,<br>no restraints) |
| ATP<sub>(Pγ)</sub>-T180<sub>(OH)</sub> | 3.76 | 3.3 / 16.5 | 10.4 / 14.4 | 13.0 | 13.5 / 5.6 |
| T180<sub>(HG1)</sub>-ATP<sub>(OG1)</sub> | 1.81 | 1.6 / 15.2 | 11.3 / 13.3 | 14.0 | 17.8 / 8.7 |
| T180<sub>(HG1)</sub>-ATP<sub>(OG2)</sub> |  | 2.9 / 17.5 | 9.1 / 14.8 | 15.2 | 15.2 / 8.1 |
| T180<sub>(HG1)</sub>-ATP<sub>(OG3)</sub> |  | 1.7 / 16.1 | 9.6 / 15.8 | 12.8 | 17.0 / 7.8 |
| ATP<sub>(Pγ)</sub>-Y182<sub>(OH)</sub> | 3.76 | 17.4 / 3.8 | 4.4 / 3.5 | 3.6 | 4.3 / 15.1 |
| Y182<sub>(HH)</sub>-ATP<sub>(OG1)</sub> | 1.81 | 17.3 / 1.9 | 5.0 / 1.4 | 3.7 | 7.3 / 19.1 |
| Y180<sub>(HH)</sub>-ATP<sub>(OG2)</sub> |  | 17.5 / 3.8 | 3.3 / 3.6 | 4.8 | 7.5 / 18.3 |
| Y182<sub>(HH)</sub>-ATP<sub>(OG3)</sub> |  | 16.9 / 3.7 | 2.9 / 3.9 | 2.6 | 7.5 / 17.9 |

**Table S9.** Statistical analysis of cellular assay (Fig. 3)

| **P38α** | **MKK6** | **Mean** | **se** | **Shapiro**<br>**Wilk** | **Welch t test**<br>**against**<br>**p38α+/**MKK6**<sup>DD</sup>** | **Significance**<br>**(<0.5 \*; <0.01**<br>**\*\*;**<br>**< 0.001 \*\*\*)** |
| --- | --- | --- | --- | --- | --- | --- |
| + | - | 2% | 0.004 | 5.42E-01 | 2.65E-09 | *** |
| + | DD | 100% | 0.044 | 2.76E-01 | 1.00E+00 | NS |
| + | K17A | 126% | 0.104 | 8.85E-02 | 3.68E-02 | * |

**Table S10.**
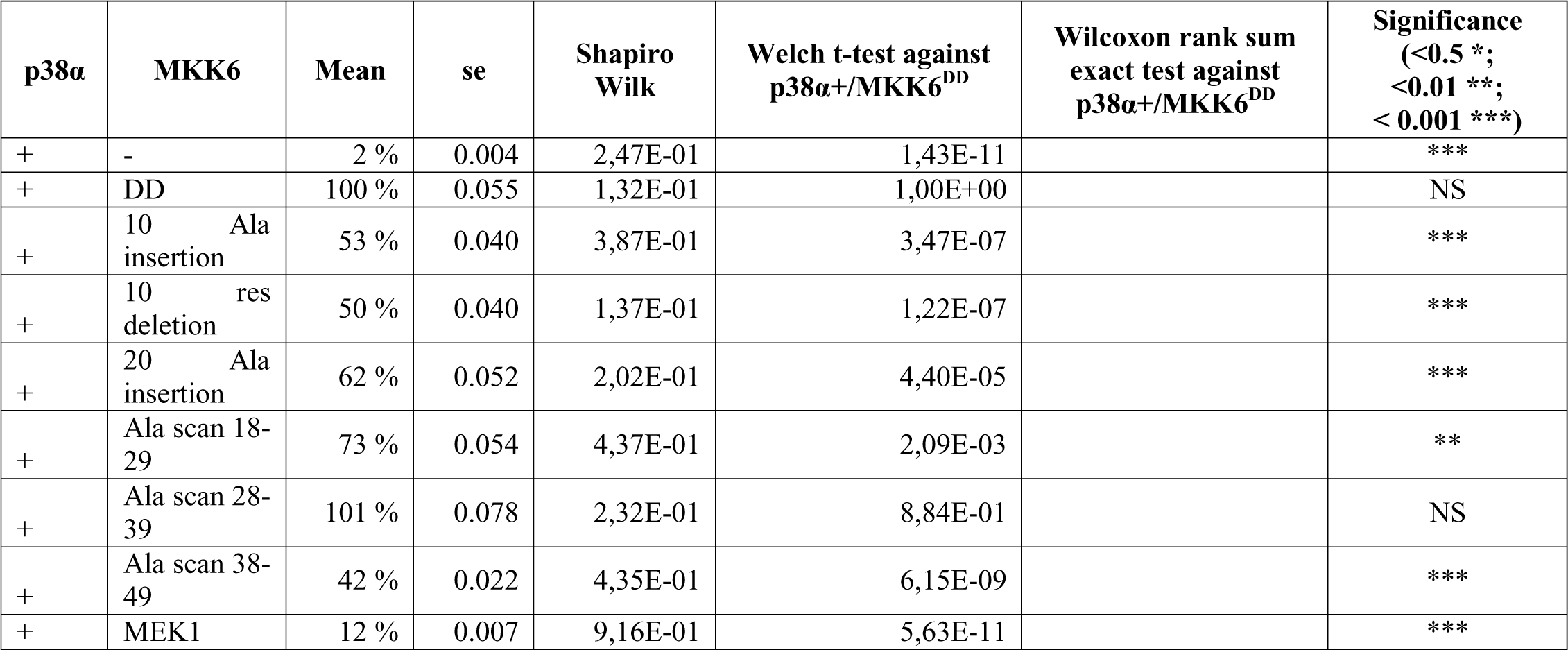
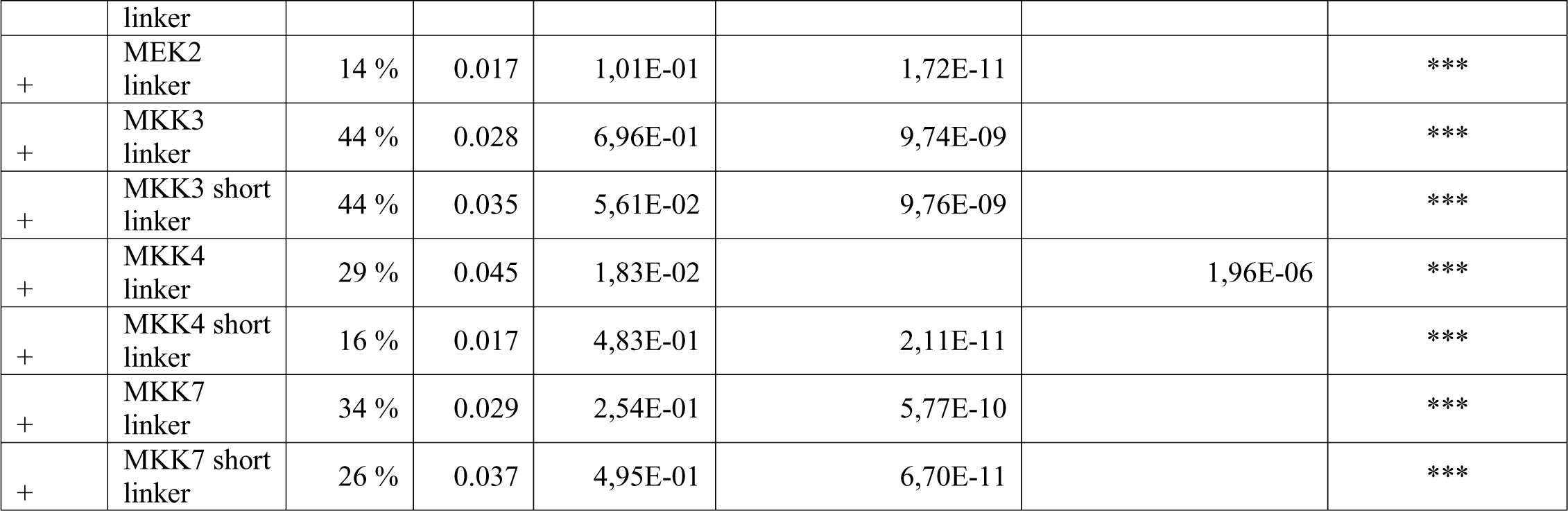
Statistical analysis of cellular assay (Fig. 5)

**Table S11.** SAXS data collection and analysis.

| Data-collection parameters |  |  |  |
| --- | --- | --- | --- |
| Instrument: | Diamond light source B21 |  |  |
| Wavelength (Å) | 0.9524 |  |  |
| q-range (Å<sup>-1</sup>) | 0.0026-0.34 |  |  |
| Sample-to-detector distance (m) | 3.7 |  |  |
| Concentration (mg/mL) | 5 |  |  |
| Temperature (K) | 277 |  |  |
| Detector | Eiger 4M (Dectris) |  |  |
| Flux (photons/s) | 2*10<sup>12</sup> |  |  |
| Beam size (µm) | 1102*240 |  |  |
| **Structural parameters**<br>**(AMP-PCP)** | **MKK6<sup>DD</sup>GRA**<br>**+ p38α<sup>WT</sup>**<br>**+ AMP-PCP** | **MKK6<sup>DD</sup>GRA**<br>**+ p38α<sup>T180V</sup>**<br>**+ AMP-PCP** | **MKK6<sup>DD</sup>GRA**<br>**+ p38α<sup>Y182F</sup>**<br>**+ AMP-PCP** |
| I0 (kDa) [from Guinier] | 3,00E-01 | 2,30E-01 | 3,00E-01 |
| Rg (Å) [from Guinier] | 41,6 | 39,1 | 39,4 |
| qminRg – qmaxRg used for Guinier | 1.4*10<sup>-2</sup>-3.1*10<sup>-2</sup> | 1.8*10<sup>-2</sup>-3.3*10<sup>-2</sup> | 1.6*10<sup>-2</sup>-3.2*10<sup>-2</sup> |
| Volume (Å<sup>3</sup>) | 1.5*10<sup>5</sup> | 1.5*10<sup>5</sup> | 1.4*10<sup>5</sup> |
| D<sub>max</sub> (Å) | 139 | 140 | 139 |
| Calculated MW | 68 | 77 | 70 |
| **Structural parameters**<br>**(ADP + AlF<sub>4</sub><sup>-</sup>)** | **MKK6<sup>DD</sup>GRA**<br>**+ p38α<sup>WT</sup>**<br>**+ ADP + AlF<sub>4</sub><sup>-</sup>** | **MKK6<sup>DD</sup>GRA**<br>**+ p38α<sup>T180V</sup>**<br>**+ ADP + AlF<sub>4</sub><sup>-</sup>** | **MKK6<sup>DD</sup>GRA**<br>**+ p38α<sup>Y182F</sup>**<br>**+ ADP + AlF<sub>4</sub><sup>-</sup>** |
| I0 (kDa) [from Guinier] | 2,60E-01 | 2,30E-01 | 2,80E-01 |
| Rg (Å) [from Guinier] | 32,2 | 29,6 | 35,3 |
| qminRg – qmaxRg used for Guinier | 1.4*10<sup>-2</sup>-4.0*10<sup>-2</sup> | 1.7*10<sup>-2</sup>-4.3*10<sup>-2</sup> | 1.4*10<sup>-2</sup>-3.6*10<sup>-2</sup> |
| Volume (Å<sup>3</sup>) | 1.3*10<sup>5</sup> | 1.2*10<sup>5</sup> | 1.4*10<sup>5</sup> |
| D<sub>max</sub> (Å) | 121 | 115 | 134 |
| Calculated MW | 65 | 59 | 69 |
| Software employed |  |  |  |
| Primary data reduction: | B21 autoprocessing pipeline |  |  |
| Data processing | Scatter IV |  |  |
| χ<sup>2</sup> determination | FoXs |  |  |

## References and Notes

1. B. Canovas, A. R. Nebreda. “Diversity and versatility of p38 kinase signalling in health and disease”, Nat Rev Mol Cell Biol 22, 346 (2021).10.1038/s41580-020-00322-w

2. M. Bouhaddou et al. “The Global Phosphorylation Landscape of SARS-CoV-2 Infection”, Cell 182, 685 (2020).10.1016/j.cell.2020.06.034

3. Daniella S. Battagello et al. “Unpuzzling COVID-19: tissue-related signaling pathways associated with SARS-CoV-2 infection and transmission”, Clinical Science 134, 2137 (2020).10.1042/cs20200904

4. S. Bellon, M. J. Fitzgibbon, T. Fox, H.-M. Hsiao, K. P. Wilson. “The structure of phosphorylated P38γ is monomeric and reveals a conserved activation-loop conformation”, Structure 7, 1057 (1999).https://doi.org/10.1016/S0969-2126(99)80173-7

5. S. S. Taylor, A. P. Kornev. “Protein kinases: evolution of dynamic regulatory proteins”, Trends in Biochemical Sciences 36, 65 (2011).https://doi.org/10.1016/j.tibs.2010.09.006

6. N. Bluthgen et al. “Effects of sequestration on signal transduction cascades”, FEBS J 273, 895 (2006).10.1111/j.1742-4658.2006.05105.x

7. F. Ortega, L. Acerenza, H. V. Westerhoff, F. Mas, M. Cascante. “Product dependence and bifunctionality compromise the ultrasensitivity of signal transduction cascades”, Proc Natl Acad Sci U S A 99, 1170 (2002).10.1073/pnas.022267399

8. M. Thattai, A. van Oudenaarden. “Attenuation of noise in ultrasensitive signaling cascades”, Biophys J 82, 2943 (2002).10.1016/S0006-3495(02)75635-X

9. D. Hammaker, G. S. Firestein. ““Go upstream, young man”: lessons learned from the p38 saga”, Annals of the Rheumatic Diseases 69, i77 (2010).10.1136/ard.2009.119479

10. K. P. Wilson et al. “Crystal structure of p38 mitogen-activated protein kinase”, J Biol Chem 271, 27696 (1996).10.1074/jbc.271.44.27696

11. X. Xie et al. “Crystal structure of JNK3: a kinase implicated in neuronal apoptosis”, Structure 6, 983 (1998).10.1016/S0969-2126(98)00100-2

12. F. Zhang, A. Strand, D. Robbins, M. H. Cobb, E. J. Goldsmith. “Atomic structure of the MAP kinase ERK2 at 2.3 Å resolution”, Nature 367, 704 (1994).10.1038/367704a0

13. X. Min et al. “The structure of the MAP2K MEK6 reveals an autoinhibitory dimer”, Structure 17, 96 (2009).10.1016/j.str.2008.11.007

14. T. Matsumoto et al. “Crystal structure of non-phosphorylated MAP2K6 in a putative auto-inhibition state”, J Biochem 151, 541 (2012).10.1093/jb/mvs023

15. J. F. Ohren et al. “Structures of human MAP kinase kinase 1 (MEK1) and MEK2 describe novel noncompetitive kinase inhibition”, Nat Struct Mol Biol 11, 1192 (2004).10.1038/nsmb859

16. M. Schroder et al. “Catalytic Domain Plasticity of MKK7 Reveals Structural Mechanisms of Allosteric Activation and Diverse Targeting Opportunities”, Cell Chem Biol 27, 1285 (2020).10.1016/j.chembiol.2020.07.014

17. Á. Garai et al. “Specificity of Linear Motifs That Bind to a Common Mitogen-Activated Protein Kinase Docking Groove”, Science Signaling 5, ra74 (2012).doi:10.1126/scisignal.2003004

18. T. Tanoue, M. Adachi, T. Moriguchi, E. Nishida. “A conserved docking motif in MAP kinases common to substrates, activators and regulators”, Nature Cell Biology 2, 110 (2000).10.1038/35000065

19. W. Peti, R. Page. “Molecular basis of MAP kinase regulation”, Protein Science 22, 1698 (2013).https://doi.org/10.1002/pro.2374

20. A. Zeke et al. “Systematic discovery of linear binding motifs targeting an ancient protein interaction surface on MAP kinases”, Mol Syst Biol 11, 837 (2015).10.15252/msb.20156269

21. T. Zhou, L. Sun, J. Humphreys, E. J. Goldsmith. “Docking interactions induce exposure of activation loop in the MAP kinase ERK2”, Structure 14, 1011 (2006).10.1016/j.str.2006.04.006

22. C. I. Chang, B. E. Xu, R. Akella, M. H. Cobb, E. J. Goldsmith. “Crystal structures of MAP kinase p38 complexed to the docking sites on its nuclear substrate MEF2A and activator MKK3b”, Mol Cell 9, 1241 (2002).10.1016/s1097-2765(02)00525-7

23. M. Dyla, M. Kjaergaard. “Intrinsically disordered linkers control tethered kinases via effective concentration”, Proc Natl Acad Sci U S A 117, 21413 (2020).10.1073/pnas.2006382117

24. J. Beenstock, N. Mooshayef, D. Engelberg. “How Do Protein Kinases Take a Selfie (Autophosphorylate)?”, Trends in Biochemical Sciences 41, 938 (2016).10.1016/j.tibs.2016.08.006

25. E. Park et al. “Architecture of autoinhibited and active BRAF-MEK1-14-3-3 complexes”, Nature 575, 545 (2019).10.1038/s41586-019-1660-y

26. Y. Kondo et al. “Cryo-EM structure of a dimeric B-Raf:14-3-3 complex reveals asymmetry in the active sites of B-Raf kinases”, Science 366, 109 (2019).10.1126/science.aay0543

27. D. F. Brennan et al. “A Raf-induced allosteric transition of KSR stimulates phosphorylation of MEK”, Nature 472, 366 (2011).10.1038/nature09860

28. P. Sok et al. “MAP Kinase-Mediated Activation of RSK1 and MK2 Substrate Kinases”, Structure 28, 1101 (2020).https://doi.org/10.1016/j.str.2020.06.007

29. E. t. Haar, P. Prabakhar, X. Liu, C. Lepre. “Crystal Structure of the P38α-MAPKAP Kinase 2 Heterodimer*”, Journal of Biological Chemistry 282, 9733 (2007).https://doi.org/10.1074/jbc.M611165200

30. L. Braun et al. “A Toxoplasma dense granule protein, GRA24, modulates the early immune response to infection by promoting a direct and sustained host p38 MAPK activation”, J Exp Med 210, 2071 (2013).10.1084/jem.20130103

31. E. Pellegrini et al. “Structural Basis for the Subversion of MAP Kinase Signaling by an Intrinsically Disordered Parasite Secreted Agonist”, Structure 25, 16 (2017).10.1016/j.str.2016.10.011

32. V. Modi, R. L. Dunbrack, Jr. “Defining a new nomenclature for the structures of active and inactive kinases”, Proc Natl Acad Sci U S A 116, 6818 (2019).10.1073/pnas.1814279116

33. R. Evans, et al. “Protein complex prediction with AlphaFold-Multimer”, bioRxiv, 2021.10.04.463034 (2022).10.1101/2021.10.04.463034

34. R. Diskin, D. Engelberg, O. Livnah. “A novel lipid binding site formed by the MAP kinase insert in p38 alpha”, J Mol Biol 375, 70 (2008).10.1016/j.jmb.2007.09.002

35. N. Tzarum, Y. Eisenberg-Domovich, J. J. Gills, P. A. Dennis, O. Livnah. “Lipid molecules induce p38alpha activation via a novel molecular switch”, J Mol Biol 424, 339 (2012).10.1016/j.jmb.2012.10.007

36. M. Buhrmann, J. Hardick, J. Weisner, L. Quambusch, D. Rauh. “Covalent Lipid Pocket Ligands Targeting p38alpha MAPK Mutants”, Angew Chem Int Ed Engl 56, 13232 (2017).10.1002/anie.201706345

37. M. Buhrmann et al. “Structure-based design, synthesis and crystallization of 2-arylquinazolines as lipid pocket ligands of p38alpha MAPK”, PLoS One 12, e0184627 (2017).10.1371/journal.pone.0184627

38. K. M. Comess et al. “Discovery and characterization of non-ATP site inhibitors of the mitogen activated protein (MAP) kinases”, ACS Chem Biol 6, 234 (2011).10.1021/cb1002619

39. O. Laufkotter, H. Hu, F. Miljkovic, J. Bajorath. “Structure- and Similarity-Based Survey of Allosteric Kinase Inhibitors, Activators, and Closely Related Compounds”, J Med Chem 65, 922 (2022).10.1021/acs.jmedchem.0c02076

40. X. H. Zhang et al. “Targeting the non-ATP-binding pocket of the MAP kinase p38γ mediates a novel mechanism of cytotoxicity in cutaneous T-cell lymphoma (CTCL)”, FEBS Letters 595, 2570 (2021).https://doi.org/10.1002/1873-3468.14186

41. K. Kirsch et al. “Co-regulation of the transcription controlling ATF2 phosphoswitch by JNK and p38”, Nat Commun 11, 5769 (2020).10.1038/s41467-020-19582-3

42. X. Liu et al. “A conserved motif in JNK/p38-specific MAPK phosphatases as a determinant for JNK1 recognition and inactivation”, Nat Commun 7, 10879 (2016).10.1038/ncomms10879

43. E. I. James, T. A. Murphree, C. Vorauer, J. R. Engen, M. Guttman. “Advances in Hydrogen/Deuterium Exchange Mass Spectrometry and the Pursuit of Challenging Biological Systems”, Chem Rev 122, 7562 (2022).10.1021/acs.chemrev.1c00279

44. S. A. Foster et al. “Activation Mechanism of Oncogenic Deletion Mutations in BRAF, EGFR, and HER2”, Cancer Cell 29, 477 (2016).https://doi.org/10.1016/j.ccell.2016.02.010

45. B. Zhang et al. “Oncogenic mutations within the β3-αC loop of EGFR/ERBB2/BRAF/MAP2K1 predict response to therapies”, Mol Genet Genomic Med 8, e1395 (2020).10.1002/mgg3.1395

46. A. Perez-Gallegos, M. Garcia-Viloca, A. Gonzalez-Lafont, J. M. Lluch. “A QM/MM study of Kemptide phosphorylation catalyzed by protein kinase A. The role of Asp166 as a general acid/base catalyst”, Phys Chem Chem Phys 17, 3497 (2015).10.1039/c4cp03579h

47. A. Kuzmanic et al. “Changes in the free-energy landscape of p38alpha MAP kinase through its canonical activation and binding events as studied by enhanced molecular dynamics simulations”, Elife 6 (2017).10.7554/eLife.22175

48. S. Doerr, G. De Fabritiis. “On-the-Fly Learning and Sampling of Ligand Binding by High-Throughput Molecular Simulations”, J Chem Theory Comput 10, 2064 (2014).10.1021/ct400919u

49. N. Plattner, S. Doerr, G. De Fabritiis, F. Noe. “Complete protein-protein association kinetics in atomic detail revealed by molecular dynamics simulations and Markov modelling”, Nat Chem 9, 1005 (2017).10.1038/nchem.2785

50. F. Pesce, K. Lindorff-Larsen. “Refining conformational ensembles of flexible proteins against small-angle x-ray scattering data”, Biophys J 120, 5124 (2021).10.1016/j.bpj.2021.10.003

51. A. H. Larsen et al. “Combining molecular dynamics simulations with small-angle X-ray and neutron scattering data to study multi-domain proteins in solution”, PLoS Comput Biol 16, e1007870 (2020).10.1371/journal.pcbi.1007870

52. S. Orioli, A. H. Larsen, S. Bottaro, K. Lindorff-Larsen. “How to learn from inconsistencies: Integrating molecular simulations with experimental data”, Prog Mol Biol Transl Sci 170, 123 (2020).10.1016/bs.pmbts.2019.12.006

53. N. S. González-Foutel et al. “Conformational buffering underlies functional selection in intrinsically disordered protein regions”, Nature Structural & Molecular Biology 29, 781 (2022).10.1038/s41594-022-00811-w

54. Y. FLEMING, et al. “Synergistic activation of stress-activated protein kinase 1/c-Jun N-terminal kinase (SAPK1/JNK) isoforms by mitogen-activated protein kinase kinase 4 (MKK4) and MKK7”, Biochemical Journal 352, 145 (2000).10.1042/bj3520145

55. C. Tournier, A. J. Whitmarsh, J. Cavanagh, T. Barrett, R. J. Davis. “The MKK7 Gene Encodes a Group of c-Jun NH_2_-Terminal Kinase Kinases”, Molecular and Cellular Biology 19, 1569 (1999).doi:10.1128/MCB.19.2.1569

56. T. O. Fischmann et al. “Crystal structures of MEK1 binary and ternary complexes with nucleotides and inhibitors”, Biochemistry 48, 2661 (2009).10.1021/bi801898e

57. V. Modi, R. L. Dunbrack. “A Structurally-Validated Multiple Sequence Alignment of 497 Human Protein Kinase Domains”, Scientific Reports 9, 19790 (2019).10.1038/s41598-019-56499-4

58. Y. Y. Zhang, J. W. Wu, Z. X. Wang. “A distinct interaction mode revealed by the crystal structure of the kinase p38α with the MAPK binding domain of the phosphatase MKP5”, Sci Signal 4, ra88 (2011).10.1126/scisignal.2002241

59. C. Dominguez, D. A. Powers, N. Tamayo. “p38 MAP kinase inhibitors: many are made, but few are chosen”, Curr Opin Drug Discov Devel 8, 421 (2005)

60. Y. L. Wang et al. “Kinetic and mechanistic studies of p38alpha MAP kinase phosphorylation by MKK6”, FEBS J 286, 1030 (2019).10.1111/febs.14762

61. J. M. Humphreys, A. T. Piala, R. Akella, H. He, E. J. Goldsmith. “Precisely Ordered Phosphorylation Reactions in the p38 Mitogen-activated Protein (MAP) Kinase Cascade*”, Journal of Biological Chemistry 288, 23322 (2013).https://doi.org/10.1074/jbc.M113.462101

62. K. Aoki, K. Takahashi, K. Kaizu, M. Matsuda. “A Quantitative Model of ERK MAP Kinase Phosphorylation in Crowded Media”, Scientific Reports 3, 1541 (2013).10.1038/srep01541

63. C. Salazar, T. Höfer. “Multisite protein phosphorylation – from molecular mechanisms to kinetic models”, The FEBS Journal 276, 3177 (2009).https://doi.org/10.1111/j.1742-4658.2009.07027.x

64. S. K. Albanese et al. “An Open Library of Human Kinase Domain Constructs for Automated Bacterial Expression”, Biochemistry 57, 4675 (2018).10.1021/acs.biochem.7b01081

65. Y. Nie, I. Bellon-Echeverria, S. Trowitzsch, C. Bieniossek, I. Berger. “Multiprotein complex production in insect cells by using polyproteins”, Methods Mol Biol 1091, 131 (2014).10.1007/978-1-62703-691-7_8

66. M. D. Tully, N. Tarbouriech, R. P. Rambo, S. Hutin. “Analysis of SEC-SAXS data via EFA deconvolution and Scatter”, J Vis Exp (2021).10.3791/61578

67. D. Schneidman-Duhovny, M. Hammel, J. A. Tainer, A. Sali. “Accurate SAXS profile computation and its assessment by contrast variation experiments”, Biophys J 105, 962 (2013).10.1016/j.bpj.2013.07.020

68. M. Schorb, I. Haberbosch, W. J. H. Hagen, Y. Schwab, D. N. Mastronarde. “Software tools for automated transmission electron microscopy”, Nat Methods 16, 471 (2019).10.1038/s41592-019-0396-9

69. A. Punjani, J. L. Rubinstein, D. J. Fleet, M. A. Brubaker. “cryoSPARC: algorithms for rapid unsupervised cryo-EM structure determination”, Nat Methods 14, 290 (2017).10.1038/nmeth.4169

70. T. Bepler et al. “Positive-unlabeled convolutional neural networks for particle picking in cryo-electron micrographs”, Nat Methods 16, 1153 (2019).10.1038/s41592-019-0575-8

71. A. J. Jakobi, M. Wilmanns, C. Sachse. “Model-based local density sharpening of cryo-EM maps”, eLife 6, e27131 (2017).10.7554/eLife.27131

72. A. Waterhouse et al. “SWISS-MODEL: homology modelling of protein structures and complexes”, Nucleic Acids Res 46, W296 (2018).10.1093/nar/gky427

73. P. V. Afonine et al. “Real-space refinement in PHENIX for cryo-EM and crystallography”, Acta Crystallogr D Struct Biol 74, 531 (2018).10.1107/S2059798318006551

74. P. Emsley, K. Cowtan. “Coot: model-building tools for molecular graphics”, Acta Crystallogr D Biol Crystallogr 60, 2126 (2004).10.1107/S0907444904019158

75. E. F. Pettersen et al. “UCSF ChimeraX: Structure visualization for researchers, educators, and developers”, Protein Sci 30, 70 (2021).10.1002/pro.3943

76. E. Pellegrini, D. Piano, M. W. Bowler. “Direct cryocooling of naked crystals: are cryoprotection agents always necessary?”, Acta Crystallogr D Biol Crystallogr 67, 902 (2011).10.1107/S0907444911031210

77. W. Kabsch. “Xds”, Acta crystallographica. Section D, Biological crystallography 66, 125 (2010).10.1107/S0907444909047337

78. M. D. Winn et al. “Overview of the CCP4 suite and current developments”, Acta crystallographica. Section D, Biological crystallography 67, 235 (2011).10.1107/S0907444910045749

79. A. Vagin, A. Teplyakov. “Molecular replacement with MOLREP”, Acta Crystallogr D Biol Crystallogr 66, 22 (2010).10.1107/S0907444909042589

80. P. V. Afonine et al. “Towards automated crystallographic structure refinement with phenix.refine”, Acta crystallographica. Section D, Biological crystallography 68, 352 (2012).10.1107/S0907444912001308

81. V. B. Chen et al. “MolProbity: all-atom structure validation for macromolecular crystallography”, Acta crystallographica. Section D, Biological crystallography 66, 12 (2010).10.1107/S0907444909042073

82. M. Mirdita et al. “ColabFold: making protein folding accessible to all”, Nature Methods 19, 679 (2022).10.1038/s41592-022-01488-1

83. E. F. Pettersen et al. “UCSF ChimeraX: Structure visualization for researchers, educators, and developers”, Protein Science 30, 70 (2021).https://doi.org/10.1002/pro.3943</otherinfo>

84. H. Wang et al. “Autoregulation of Class II Alpha PI3K Activity by Its Lipid-Binding PX-C2 Domain Module”, Mol Cell 71, 343 (2018).10.1016/j.molcel.2018.06.042

85. Y. Perez-Riverol et al. “The PRIDE database resources in 2022: a hub for mass spectrometry-based proteomics evidences”, Nucleic Acids Res 50, D543 (2022).10.1093/nar/gkab1038

86. C. S. Hughes et al. “Ultrasensitive proteome analysis using paramagnetic bead technology”, Mol Syst Biol 10, 757 (2014).10.15252/msb.20145625

87. H. Franken et al. “Thermal proteome profiling for unbiased identification of direct and indirect drug targets using multiplexed quantitative mass spectrometry”, Nat Protoc 10, 1567 (2015).10.1038/nprot.2015.101

88. R. A. Friesner et al. “Extra precision glide: docking and scoring incorporating a model of hydrophobic enclosure for protein-ligand complexes”, J Med Chem 49, 6177 (2006).10.1021/jm051256o

89. A. Sali, T. L. Blundell. “Comparative protein modelling by satisfaction of spatial restraints”, J Mol Biol 234, 779 (1993).10.1006/jmbi.1993.1626

90. G. Martinez-Rosell, T. Giorgino, G. De Fabritiis. “PlayMolecule ProteinPrepare: A Web Application for Protein Preparation for Molecular Dynamics Simulations”, J Chem Inf Model 57, 1511 (2017).10.1021/acs.jcim.7b00190

91. K. L. Meagher, L. T. Redman, H. A. Carlson. “Development of polyphosphate parameters for use with the AMBER force field”, J Comput Chem 24, 1016 (2003).10.1002/jcc.10262

92. F. P. Buelens, H. Leonov, B. L. de Groot, H. Grubmuller. “ATP-Magnesium Coordination: Protein Structure-Based Force Field Evaluation and Corrections”, J Chem Theory Comput 17, 1922 (2021).10.1021/acs.jctc.0c01205

93. M. J. Abraham et al. “GROMACS: High performance molecular simulations through multi-level parallelism from laptops to supercomputers”, SoftwareX 1–2, 19 (2015).https://doi.org/10.1016/j.softx.2015.06.001

94. G. A. Tribello, M. Bonomi, D. Branduardi, C. Camilloni, G. Bussi. “PLUMED 2: New feathers for an old bird”, Computer Physics Communications 185, 604 (2014).https://doi.org/10.1016/j.cpc.2013.09.018

95. G. Bussi, D. Donadio, M. Parrinello. “Canonical sampling through velocity rescaling”, The Journal of Chemical Physics 126, 014101 (2007).10.1063/1.2408420

96. M. Bernetti, G. Bussi. “Pressure control using stochastic cell rescaling”, The Journal of Chemical Physics 153, 114107 (2020).10.1063/5.0020514

97. P. consortium. “Promoting transparency and reproducibility in enhanced molecular simulations”, Nat Methods 16, 670 (2019).10.1038/s41592-019-0506-8

98. M. Marchi, P. Ballone. “Adiabatic bias molecular dynamics: A method to navigate the conformational space of complex molecular systems”, The Journal of Chemical Physics 110, 3697 (1999).10.1063/1.478259

99. S. Doerr, M. J. Harvey, F. Noe, G. De Fabritiis. “HTMD: High-Throughput Molecular Dynamics for Molecular Discovery”, J Chem Theory Comput 12, 1845 (2016).10.1021/acs.jctc.6b00049

100. G. Pérez-Hernández, F. Paul, T. Giorgino, G. D. Fabritiis, F. Noé. “Identification of slow molecular order parameters for Markov model construction”, The Journal of Chemical Physics 139, 015102 (2013).10.1063/1.4811489

101. P. Raiteri, A. Laio, F. L. Gervasio, C. Micheletti, M. Parrinello. “Efficient reconstruction of complex free energy landscapes by multiple walkers metadynamics”, J Phys Chem B 110, 3533 (2006).10.1021/jp054359r

102. X. Daura et al. “Peptide Folding: When Simulation Meets Experiment”, Angewandte Chemie International Edition 38, 236 (1999).https://doi.org/10.1002/(SICI)1521-3773(19990115)38:1/2<236::AID-ANIE236>3.0.CO;2-M

103. S. Röblitz, M. Weber. “Fuzzy spectral clustering by PCCA+: application to Markov state models and data classification”, Advances in Data Analysis and Classification 7, 147 (2013).10.1007/s11634-013-0134-6

104. S. Bottaro, G. Bussi, S. D. Kennedy, D. H. Turner, K. Lindorff-Larsen. “Conformational ensembles of RNA oligonucleotides from integrating NMR and molecular simulations”, Sci Adv 4, eaar8521 (2018).10.1126/sciadv.aar8521

105. S. Grudinin, M. Garkavenko, A. Kazennov. “Pepsi-SAXS: an adaptive method for rapid and accurate computation of small-angle X-ray scattering profiles”, Acta Crystallographica Section D 73, 449 (2017).doi:10.1107/S2059798317005745

106. I. Galdadas et al. “Structural basis of the effect of activating mutations on the EGF receptor”, eLife 10, e65824 (2021).10.7554/eLife.65824

107. Y. Shan, A. Arkhipov, E. T. Kim, A. C. Pan, D. E. Shaw. “Transitions to catalytically inactive conformations in EGFR kinase”, Proceedings of the National Academy of Sciences 110, 7270 (2013).doi:10.1073/pnas.1220843110

108. J. Eswaran et al. “Structure of the Human Protein Kinase MPSK1 Reveals an Atypical Activation Loop Architecture”, Structure 16, 115 (2008).https://doi.org/10.1016/j.str.2007.10.026

